# Inhibition of Gyrase via a conserved α-hairpin in *Vibrio cholerae* ParE2 and neutralization by ParD2

**DOI:** 10.1101/2025.07.24.666505

**Authors:** Yana Girardin, Remy Loris

## Abstract

The ParE family of toxins is known to target Gyrase through an as yet unknown molecular mechanism. Here we show that *Vibrio cholerae* ParE2 (*Vc*ParE2) interacts with Gyrase in a DNA-dependent manner. The interaction site is located at the ParE-specific α-hairpin and involves Trp25 as a key residue. The latter is poorly conserved within the ParE family, although full toxicity is only retained upon substitution with Tyr in the *Vc*ParE2 context. *In vitro*, the Trp25Ala mutation reduces the rate of Gyrase-mediated supercoiling, while the so-called cleavable complex remains stabilized. The C-terminal domain of the antitoxin *Vc*ParD2, which wraps its intrinsically disordered domain around *Vc*ParE2 without covering the Trp25-containing interaction site, binds to *Vc*ParE2 with high affinity, but only partially prevents ParE-mediated growth inhibition. Full inhibition of *Vc*ParE2 requires full length *Vc*ParD2 that leads to a complex where Trp25 is fully shielded from solvent.

## INTRODUCTION

In 1928, Alexander Fleming discovered the first true antibiotic, penicillin [1]. This discovery was the beginning of a cascade of identifying several antibiotic classes, marking the ‘golden age of antibiotics’ between the 1940s and 1960s. The majority of these antibiotics remain in use today, however their effectiveness is increasingly compromised by the rise of antibiotic resistance (for a review, see [2]).

An important target of different classes of antibiotics is Gyrase, as it is not present in Eukaryotes. This enzyme is a heterotetrameric type II topoisomerase involved in crucial processes, such as DNA replication and transcription. Unlike other topoisomerases, it relaxes positive supercoiled DNA and is capable of introducing negative supercoils at the expense of ATP hydrolysis [3]. Without Gyrase activity, overwinding and entanglement would occur during replication and transcription, leading to cell death. Gyrase consists of two 97 kDa GyrA subunits and two 90 kDa GyrB subunits [4]. GyrB contains an N-terminal ATPase domain and a topoisomerase-primase domain at the C-terminus. GyrA has an N-terminal breakage-reunion domain, composed of a winged-helix domain that houses the primary catalytic residue tyrosine 122, as well as a tower domain and a coiled-coil domain. Its C-terminal domain contains a β-pinwheel fold and is responsible for DNA binding and wrapping [5–8].

In addition to antibiotics, several toxins from toxin-antitoxin (TA) systems (namely CcdB, ParE, FicT and TsbT) have been shown to target Gyrase [9–12]. Toxin-antitoxin systems are two-component systems encoded on a small two-gene operon and consist of an inhibitor of cell growth (the toxin) and a neutralizing regulator (the antitoxin). The toxin targets specific cellular functions, inhibiting cell growth and in some cases causing cell death, while the antitoxin neutralizes the toxin’s activity. TA systems are widely distributed on prokaryotic plasmids and chromosomes [13], where they participate in various important processes, including plasmid stabilisation [14–15], phage inhibition [16–19], biofilm formation [20] and persistence [21]; for a review see [22]. Currently, eight general types of TA systems have been characterized, based on the nature and mode of action of the antitoxin (for a review see [23–24]). Among these, the Type II TA systems are the most common and best studied. Here, both toxin and antitoxin are proteins, and the antitoxin inhibits the toxin via the formation of a non-covalent complex.

Type II TA systems include many otherwise unrelated families, one of them being the parDE family. The latter was initially discovered on *Escherichia coli* plasmid RK2 [25], but since then many homologs have been identified on different bacterial chromosomes, including three parDE systems (*parDE1, parDE2 and parDE3*) located on the small chromosome of Vibrio cholerae [26]. When investigated, ParE toxins - including *Vc*ParE2 from *V. cholerae* - were shown to inhibit DNA Gyrase through a yet unknown molecular mechanism that appears different from that of the well-studied inhibitors CcdB and quinolones [10,27]. Previous studies report inconsistent findings regarding the Gyrase regions targeted by different ParE homologues. For instance, *Vc*ParE2 was reported to target the N-terminal domain of GyrA (GyrA59) [27]. In contrast, no direct interaction with GyrA was observed for *E. coli* O157:H7 ParE [28]. Furthermore, *Mycobacterium tuberculosis* ParE2 was reported to target the GyrB subunit and *Pseudomonas aeruginosa* ParE to interact with a fused GyrB-GyrA construct [29,30]. Overall, the identification of critical residues responsible for ParE activity remains challenging to date, largely due to the lack of structural data on the ParE-Gyrase interaction and the limited sequence conservation among different members of the ParE family [31].

This paper describes a systematic mutagenesis study which identifies the Gyrase-interaction site of *Vc*ParE2 to be located at the ParE-specific α-hairpin, with Trp25 as a key residue. Via a series of *in vivo* and in *vitro* studies, further details are revealed on how *Vc*ParE2 toxicity is regulated by the antitoxin *Vc*ParD2.

## MATERIALS AND METHODS

### Protein Purification and Complex Formation

VcParE2, *E. coli* GyrA59 (a 59 kDa N-terminal fragment of GyrA), *E. coli* GyrA and *E. coli* GyrB were expressed and purified following the previously described methods [32–33,27]. Full Gyrase was obtained by combining equimolar amounts of GyrA and GyrB subunits and subsequently incubating the mixture for 30 minutes at room temperature.

### Isothermal Titration Calorimetry

Prior to the isothermal titration calorimetry measurements, all proteins were dialyzed against 20 mM Tris pH 7.5, 200 mM NaCl, 5 mM MaCl_2_, 1 mM TCEP, 1% glycerol, 0.02% Tween. The syringe of the MicroCal PEAQ system was filled with 75 µM of Gyrase A59, Gyrase A, Gyrase B or full Gyrase, while the cell was loaded with 7.5 µM *Vc*ParE2. A total of 24 injections of 1.5 μl Gyrase were titrated into the cell, resulting in a final Gyrase:VcParE2 molar ratio of 2. All titrations were performed at stirring speed 750 rpm and a constant temperature of 25°C. The collected ITC data were processed using the MicroCal PEAQ-ITC Analysis Software.

### Biolayer Interferometry

All binding experiments were performed at 25°C with a shaking speed of 1000 rpm, except during the dissociation phase, where it was increased to 1200 rpm. An unloaded control sensor was included to assess nonspecific binding.

#### VcParE2-VcParD2 fragments on Ni-NTA Biosensors

Interaction studies between *Vc*ParE2 and peptides corresponding to four C-terminal truncates of *Vc*ParD2 (98% purity, Synpeptide; Table 1) were conducted in running buffer (20 mM Tris pH 8.0, 150 mM NaCl, 0.1 mM TCEP, 0.5% BSA, 0.05% Tween) using the OctetRED96 (fortéBIO) system. Dip and read^TM^ Ni-NTA biosensors (fortéBIO) were pre-hydrated for max. 30 min. After a 60 s baseline stabilization, 10 µg/mL *Vc*ParE2 was immobilized for 90 s via its C-terminal his-tag. Unbound *Vc*ParE2 was removed by a 60 s wash. The association phase was then monitored for 400 s by exposing the sensors to *Vc*ParD2 truncate concentrations of 0.1, 1 and 5 µM, followed by a 600 s dissociation phase.

**Table 1:** List of peptides corresponding with different C-terminal truncates of *Vc*ParD2. E41Y and S76Y mutations (underlined) were introduced to facilitate determination of the peptide concentration. Both mutations are not expected to alter interaction with *Vc*ParE2, as they are pointing away from the toxin in the *Vc*ParDE2 complex (PDB ID: 7R5A).

| Peptide name | VcParD2 residues | Amino acid sequence |
| --- | --- | --- |
| VcParD2<sup>CTD</sup> | E41-R80 | NH2-YNQETKLQSLRQLLIEGEQSGDADYDLDSFINELDYENIR-COOH |
| VcParD2<sup>N-CTD</sup> | E41-S60 | NH2-YNQETKLQSLRQLLIEGEQS-COOH |
| VcParD2<sup>C-CTD</sup> | D66-R80 | NH2-DLDSFINELDYENIR-COOH |
| VcParD2<sup>LC-CTD</sup> | G61-R80 | NH2-DADYDLDSFINELDYENIR-COOH |

#### VcParE2 and his-GyrA59 on AR2G Biosensors

*Vc*ParE2 or his-GyrA59 was immobilized on pre-hydrated AR2G biosensors (Sartorius) via lysine coupling. Sensors were pre-hydrated for 30 min and activated (300 s) in 20 mM 1-ethyl-3-(3-dimethylaminopropyl)carbodiimide (EDC) and 10 mM sulfo-N-hydroxysuccinimide (NHS), then loaded with 5 µg/mL ligand in 10 mM sodium acetate, 0.05% Tween, 0.5% BSA (pH 6.0). Immobilization was quenched in 1 M ethanolamine (pH 8.5) and unbound protein was removed by a 60 s wash in 20 mM Tris pH 8.0, 500 mM NaCl, 1 mM TCEP, 0.05% Tween, 0.5% BSA. Analytes were dialyzed into this buffer before analysis. Association was measured by a 900 s incubation in 20 mM analyte (his-GyrA59 or *Vc*ParE2, respectively), followed by 900 s dissociation.

#### VcParE2 and his-GyrA59/his-GyrA on Streptavidin Biosensors

The N-terminal α-amino groups of *Vc*ParE2, his-GyrA59 and his-GyrA (in 37.7 mM Na_2_HPO_4_, 12.3 mM NaH_2_PO_4_xH_2_O pH 6.5, 500 mM NaCl, 1 mM TCEP) were biotinylated by mixing then with a five-fold molar excess of EZ-Link™ NHS-PEG4-Biotin (ThermoFisher), followed by a 24-hour incubation at 7°C. To remove unbound biotin, the protein was injected on a Superdex 75 Increase column equilibrated with phosphate buffer (37.7 mM Na_2_HPO_4_, 12.3 mM NaH_2_PO_4_xH_2_O pH 7.6, 500 mM NaCl, 1 mM TCEP). For biosensor loading, 5 µg/mL biotinylated his-GyrA59 or his-GyrA was immobilized on streptavidin biosensors (SA, Sartorius) pre-hydrated in phosphate buffer supplemented with 0.5% BSA and 0.05% Tween. After a 60 s wash, association with 2 µM *Vc*ParE2 was monitored for 1000 s, followed by a 700 s dissociation phase. The procedure was repeated for 5 µg/mL biotinylated *Vc*ParE2 submerged in 100, 50 and 25 µM his-GyrA59.

### Electrophoretic Mobility Shift Assay

A 286-bp-long DNA fragment encoding the bacteriophage Mu strong Gyrase Site (NCBI NC_000929: 17765-18050, [34]) was obtained via PCR amplification of a synthetically ordered DNA fragment using following primers: 5’-TTCCTGCGCGTCCTTATATG-3’ and 5’-GCTGCTCATCATGGTTTGTG-3’. The PCR was run under following conditions: 3 minutes initial denaturation at 94°C, 30 cycles of 30 s denaturation at 94°C, 30 s annealing at 58°C and 20 s elongation at 72°C, and a final extension of 3 minutes at 72°C. The resulting DNA fragment was cleaned with the Monarch® PCR and DNA cleanup kit. Of this DNA, 30 nM was mixed with either 3 µM Gyrase or 3 µM Gyrase and 15 µM *Vc*ParE2. Complex formation was allowed by a 30-minute incubation step at 37°C. Each sample was mixed with 6X 30% glycerol loading dye, of which 15 µL was loaded on a 1% agarose gel and run for 20 minutes at 200 V on ice, using 1X Tris-Glycine-EDTA as running buffer. GelRed® (Biotium) was used to stain the DNA.

### Production of Chemically Competent Cells

Competent *E. coli* cells were prepared using the CaCl_2_ method [35]. A dilution of 250 µL overnight preculture in 25 mL LB medium was set to incubate at 37°C until the OD_600_ reached 0.2. The cells were then cooled on ice for 15 minutes and centrifugated for 5 minutes at 4000 rpm and 4°C. Each pellet was resuspended in 10 mL of ice-cold 0.1 M CaCl_2_, cooled on ice for 25 minutes, and centrifuged as described before. The resulting pellet was resuspended in a mixture of 250 µL ice-cold 0.1 M CaCl_2_ and 50 µL 80% glycerol. Aliquots of 50 µL were flash-frozen with liquid nitrogen and stored at −80°C.

### Accessible Solvent Area Calculation

Residues to be included in the single alanine scan were identified by calculating the solvent-accessible area (SAA) of each side chain in the *Vc*ParE2 monomer. *Vc*ParE2 coordinates, extracted from the *Vc*ParDE2 complex (PDB ID: 7R5A, chain A), were imported into the web-based ‘Accessible Surface Area and Accessibility Calculation for Protein’ tool provided by the Center for Informational Biology (http://cib.cf.ocha.ac.jp/bitool/ASA/; Ochanomizu University; version 1.2). The resulting surface area contributions of all R-group atoms were summed and residues with a side chain SAA below 10.5 Å^2^ were excluded from the scan. Also, glycines, prolines, alanines and the N-terminal methionine were not taken into consideration.

### Cloning and Transformation of the *Vc*ParE2 Mutants

The pET28b expression vector was purified using the Monarch® Plasmid Miniprep Kit (New England Biolabs) and digested by incubation with NcoI-HF and HindIII-HF restriction enzymes at 37°C for 1 hour. Enzyme activity was terminated by incubation at 80°C for 20 minutes. Synthetic DNA fragments encoding the *Vc*ParE2 mutants and Small Ubiquitin-like Modifier (SUMO)-variants (Supplementary Table S1), obtained via Twist Bioscience, were dissolved to a concentration of 50 ng/µL and assembled into the digested pET28b vector using HiFi DNA assembly (New England Biolabs). Two microliters of the assembly mix were transformed into competent NEB5α cells that had been thawed on ice. The cells incubated on ice for 30 minutes, followed by a 2-minute heat shock at 42°C and a 10-minute incubation on ice. Transformed cells were then combined with 1 mL LB medium and incubated for 4h at 37°C to allow phenotypical expression. Cells were pelletized by centrifugation at 17000g for 1 minute. After removing 850 µL of supernatant, the cells were resuspended in the remaining LB medium. 100 µL of the resulting cell suspension was plated on LB agar plates containing 50 µg/mL kanamycin and incubated overnight at 37°C. Colonies obtained from the agar plate were selected using a sterile toothpick and diluted in 30 µL sterile H_2_O, with 1.5 µL used as template for colony PCR with Ex Taq DNA polymerase (Takara) and the following primers: T7.FW (5’-GCGAAATTAATACGACTCACTATAG-3’) and T7.RV (5’-GCTAGTTATTGCTCAGCGG-3’). The PCR protocol consisted of an initial denaturation step at 95°C for 3 minutes, followed by 30 cycles of denaturation at 95°C for 30 seconds, annealing at 53°C 30 seconds, and elongation at 72°C for 40 seconds. A final elongation step was performed at 72°C for 3 minutes. The resulting PCR products were analyzed by agarose gel electrophoresis to confirm their size. Correctly sized fragments were purified using the Monarch® PCR and DNA Cleanup Kit (New England Biolabs) and sequenced by Sanger Sequencing (Eurofins).

### Spot Tests

Synthetic DNA fragments (Twist Biosciences) encoding *Vc*ParE2 variants under the control of an arabinose promoter and a theophylline riboswitch or a tac promoter and a theophylline riboswitch (P_*BAD*_-RS_theo_ and Ptac-RS_theo_, [36]) were inserted in the BglII and XhoI restriction sites of the pET22b vector using Gibson assembly. The resulting constructs were subsequently transformed in *E. coli* EPI400 cells.

Confirmation of successful transformation was obtained by colony PCR and Sanger sequencing using primers 5’-GATCTTCCCCATCGGTGATG-3’ and 5’- GCAGCAGCCAACTCAGCTTC-3’. For each *Vc*ParE2 variant, a 5 mL preculture was inoculated and grown overnight at 37°C. All precultures were then adjusted to an equal OD_600_ of 2.8 and dilution series ranging from 10^0^ to 10^-6^ were prepared in LB medium. From each dilution, 5 µL was spotted on two types of 50 mL LB agar plates. A first one enriched with 100 µg/mL ampicillin, corresponding with the OFF-state, and a second one supplemented with 100 µg/mL ampicillin, 0.1% arabinose and 2 mM theophylline, corresponding with the ON-state promoting protein production (in case of the P_*BAD*_-RS_theo_ constructs). For P*tac*-RS_theo_ constructs, the ON-plate was enriched with 0.5 mM IPTG instead of arabinose. The spotted plates were incubated overnight at 37°C.

A similar protocol was followed for *E. coli* EPI400 carrying both (i) a pET22b plasmid encoding the previously described P_BAD_-RS_theo_-VcParE2 and (ii) a pET28b plasmid having one of the following genes under control of a *tac* promoter introduced at the BglII and XhoI restriction sites: *VcparD2, VcparD2*^E41-R80^, *VcparD2*^E41-S60^, *VcparD2*^D66-R80^, *VcparD2*^G61-R80^. OFF-state LB plates contained 100 µg/mL ampicillin, 50 µg/mL kanamycin and 0.5 mM IPTG. ON-state LB plates were supplemented with 100 µg/mL ampicillin, 50 µg/mL kanamycin, 0.5 mM IPTG, 2 mM theophylline and 0.1% arabinose.

### Expression and Purification of *Vc*ParE2, *Vc*ParE2W25A or *Vc*ParE2F20A,W25A

BL21 (DE3) cells were transformed with a pET28a vector containing a T7 promoter upstream of three variants of the bicistronic *par*DE2 gene: the wild-type variant encoding *Vc*ParD2 and a C-terminally histidine-tagged *Vc*ParE2; the single mutant variant where tryptophane at position 25 in *Vc*ParE2 is substituted with alanine (from hereon called *Vc*ParE2^W25A^); and double mutant variant having phenylalanine at position 20 and tryptophane at position 25 of *Vc*ParE2 substituted with alanine (from hereon called *Vc*ParE2^F20A, W25A^). The three different *Vc*ParD2-VcParE2 complexes were expressed and purified according to the protocol outlined in [37]. Complexes were separated in their building blocks (free *Vc*ParD2 and *Vc*ParE2, *Vc*ParE2^W25A^ or *Vc*ParE2^F20A, W25A^) by applying the unfolding-refolding method described in [32].

### Analytical Gel Filtration

A 500 µL sample of either *Vc*ParE2, *Vc*ParE2^W25A^ or *Vc*ParE2^F20A, W25A^ at a concentration of 0.31 mg/mL was loaded onto a Superdex increase 75 10/300 column (Cytiva) pre-equilibrated with 20 mM Tris pH 7.4, 500 mM NaCl and 1 mM TCEP. The column was calibrated under the same buffer conditions using the Bio-Rad molecular mass standard, which consist of bovine thyroglobulin (670 kDa), bovine γ-globulin (158 kDa), chicken ovalbumin (44 kDa), horse myoglobin (17 kDa) and vitamin B12 (1.35 kDa).

### Circular Dichroism

CD measurements were performed on 200 µL of *Vc*ParE2, *Vc*ParE2^W25A^ and *Vc*ParE2^F20A, W25A^ at 0.20 mg/mL in 20 mM Tris pH 7.4, 500 mM NaCl, 1 mM TCEP. Each sample was introduced in a 1 mm quartz cuvette (Hellma) and spectra were recorded in triplicate at room temperature with a MOS-500 spectropolarimeter (BioLogic), scanning wavelengths ranging from 200 to 250 nm in 1 nm steps. The buffer signal was subtracted from the average CD signal (θ in mdeg) and it was converted to molar ellipticity ([θ] in deg cm^2^ dmol^−1^) using the formula: [θ] = (θ.MM)/(10.c.l.N), where MM is the molecular weight (Da), c the protein concentration (mg/mL), l the cuvette path length (cm) and N number of amino acids.

### Gyrase and Topoisomerase IV Activity Assays

#### Supercoiling assays

For Gyrase and Topoisomerase IV activity assays were performed using the kits provided by Inspiralis.

#### Supercoiling assays

The *E. coli* Gyrase supercoiling assay was applied to 50 µM of *Vc*ParE2, *Vc*ParE2^W25A^ and *Vc*ParE2^F20A, W25A^ following the protocol described in [32].

#### Cleavage assays

Supercoiled pBR322 plasmid DNA (0.5 µg) was incubated with *Vc*ParE2, *Vc*ParE2^W25A^, *Vc*ParE2^F20A, W25A^ or moxifloxacin, each at a final concentration of 5 µM. For Gyrase cleavage assays, 5 U of Gyrase were added to the reaction mixture. The assay buffer consisted of 35 mM Tris-HCl (pH 7.5), 24 mM KCl, 4 mM MgCl₂, 2 mM DTT, 1.8 mM spermidine, 6.5% glycerol, 0.1 mg/mL albumin and 0.1 mg/mL BSA. For topoisomerase IV cleavage assays, 5 U of Topoisomerase IV were added. The assay buffer contained 40 mM HEPES-KOH (pH 7.6), 100 mM potassium glutamate, 10 mM magnesium acetate, 10 mM DTT and 50 µg/mL albumin. Reactions were conducted both in the absence and presence of 1 mM ATP. Topoisomerase-mediated DNA cleavage was performed by incubating the reaction mixture at 37°C for 1 hour. Following incubation, 0.5 mg/mL proteinase K and 0.2% SDS were added, and the mixture was incubated for an additional 30 minutes at 37°C. Reactions were terminated by adding an equal volume of GSTEB (40% glycerol, 100 mM Tris-HCl pH 8.0, 1 mM EDTA, 0.5 mg/mL bromophenol blue) and chloroform/isoamyl alcohol (24:1). Samples were vortexed, centrifuged at 10,000 × g for 1 minute and 20 µL of the aqueous phase was loaded onto a 1% agarose gel prepared in 1X TAE buffer. Electrophoresis was carried out in 1X TAE buffer at 80 V for 2 hours in the absence of intercalators. DNA was stained post-electrophoresis with GelRed (Biotium) and visualized using a GelDoc imaging system (Bio-Rad).

## RESULTS

### The *Vc*ParE2-Gyrase Complex is only Formed in the Context of a DNA Target

Earlier work indicated that *Vc*ParE2 interacts directly with the GyrA59 fragment of the A subunit of Gyrase [27]. Attempts were made to confirm this results, but contrary to previous surface plasmon resonance measurements, no *in vitro* interaction was observed between *Vc*ParE2 and GyrA59 using biolayer interferometry (BLI) (Supplementary Figure S1 A-D).

Using lysine coupling of either GyrA59 or *Vc*ParE2 for immobilization, only non-specific binding occurs upon submerging in the interaction partner, regardless of whether *Vc*ParE2 or GyrA59 was immobilized. Similarly, an alternative immobilization approach (biotinylation of the primary amino groups of *Vc*ParE2 and GyrA59 followed by streptavidin coupling) yields an identical outcome: no detectable interaction between *Vc*ParE2 and GyrA59, irrespective of which protein was immobilized. This was also the case when using full length GyrA instead of GyrA59. The incapability of *Vc*ParE2 to interact with Gyrase or its subunits was further confirmed by ITC measurements, where *Vc*ParE2 titration with GyrA59, GyrA, GyrB or full Gyrase failed to demonstrate binding (Supplementary Figure S1 E-H).

In contrast, *Vc*ParE2 does inhibit Gyrase-mediated supercoiling of pBR322 and stabilizes the cleavage complex in a concentration-dependent manner (Figure 1 A,B) as was also observed previously [27,32]. To evaluate whether *Vc*ParE2 would interact with Gyrase in its DNA-bound state, an EMSA experiment was performed using a 286 bp DNA fragment encoding the Strong Gyrase site located at the center of the bacteriophage Mu genome [34]. As expected, Gyrase binds to this fragment and induces a mobility shift of the DNA. Further addition of *Vc*ParE2 results in a further shift, indicating the formation of a ternary complex (Figure 1 C). This strongly suggests that either *Vc*ParE2 interacts with DNA-bound Gyrase or that Gyrase must be actively catalyzing supercoiling for the toxin to bind.

**Figure 1:**
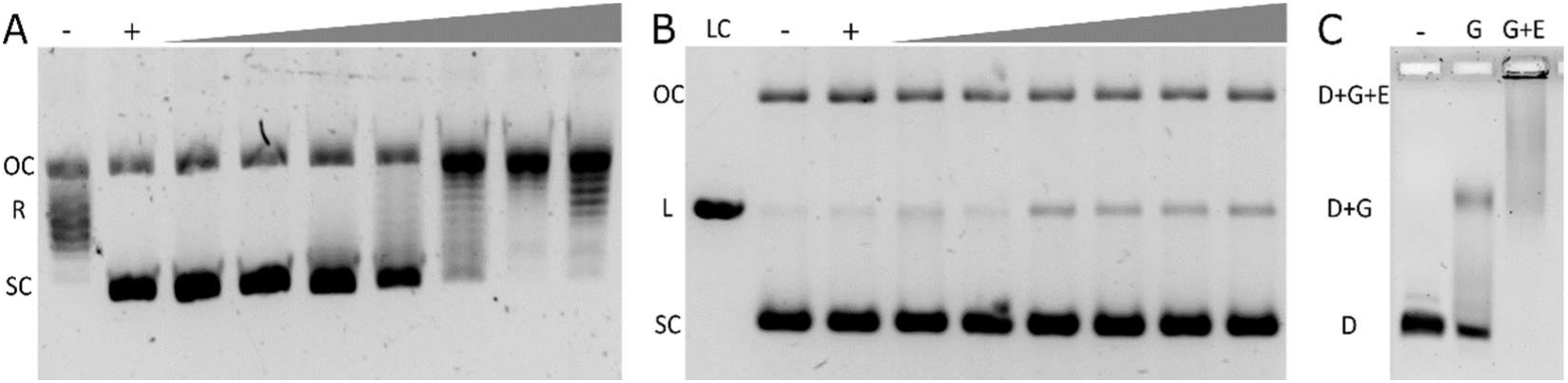
VcParE2 requires DNA for its interaction with Gyrase. (**A**) Titration of *Vc*ParE2 inhibits Gyrase-mediated supercoiling of the pBR322 plasmid. DNA topoisomers are labelled as OC (open circular), R (relaxed) and SC (supercoiled). (**B**) Increasing concentrations of *Vc*ParE2 lead to stabilisation of the Gyrase-DNA cleavage complex. DNA topoisomers are labelled as OC (open circular), L (linear) and SC (supercoiled). (**C**) EMSA showing titration of Gyrase alone (**G**) and Gyrase with *Vc*ParE2 (G+E) into a 286 bp DNA fragment, resulting in a mobility shift of the DNA. The negative control (-) shows relaxed plasmid in absence of Gyrase, while the positive control (+) represents negatively supercoiled plasmid due to addition of Gyrase.

### Tryptophane 25 is Essential for *Vc*ParE2 Activity

To pinpoint residues crucial for *Vc*ParE2 activity, an alanine scan was performed targeting all surface-exposed *Vc*ParE2 residues with a side chain solvent accessibility exceeding 10 Å (Supplementary Table S2). Synthetically designed DNA fragments of each alanine mutant were inserted at the NcoI and HindIII restriction sites of a pET28b vector and colony formation was evaluated after transformation in *E. coli* NEB5α cells. Due to leaky expression, no cell growth was observed for wild-type nor for 70 out of 71 screened single alanine mutants without accumulation of additional frameshift mutations (Supplementary Table S2, Supplementary Figure S2). Only the variant harboring the mutation Trp25Ala (from hereon called *Vc*ParE2^W25A^) is viable, although still only very few colonies are observed (Figure 2). This indicates that although the Trp25Ala mutant is less active than WT, it still shows significant toxicity.

**Figure 2:**
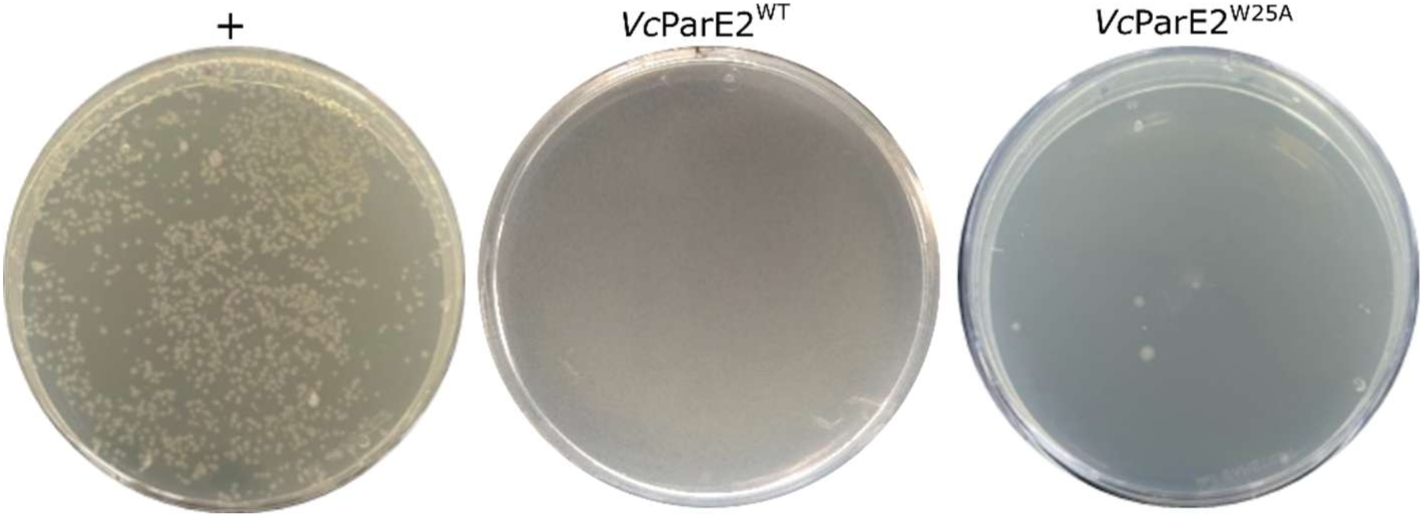
LB kanamycin plates after transformation of NEB5α cells with empty pET28b vector (+), wild-type *Vc*ParE2 and *Vc*ParE2^W25A^.

To further quantify the toxicity of wild-type *Vc*ParE2 and *Vc*ParE2^W25A^, the previously developed P_BAD_-RS_theo_ multi-layer control system for cloning highly toxic proteins was used [36]. Here, the genes encoding *Vc*ParE2 mutants are introduced in a pET22b vector under control of an inducible P_BAD_ promoter and a theophylline riboswitch. Spot tests show that without induction, cells encoding the wild-type protein grow efficiently, as do the ones that only carry an empty vector (Figure 3). Upon induction, however, cells expressing wild-type *Vc*ParE2 show a 10^4^-fold growth reduction, while cells harboring the empty vector remain unaffected. The *Vc*ParE2^W25A^ mutant exhibits an intermediate behavior where viability is strongly reduced, but not completely abolished. This indicates that Trp25 plays an important role in *Vc*ParE2 activity but does not on its own account for the full extent of its toxicity.

**Figure 3:**
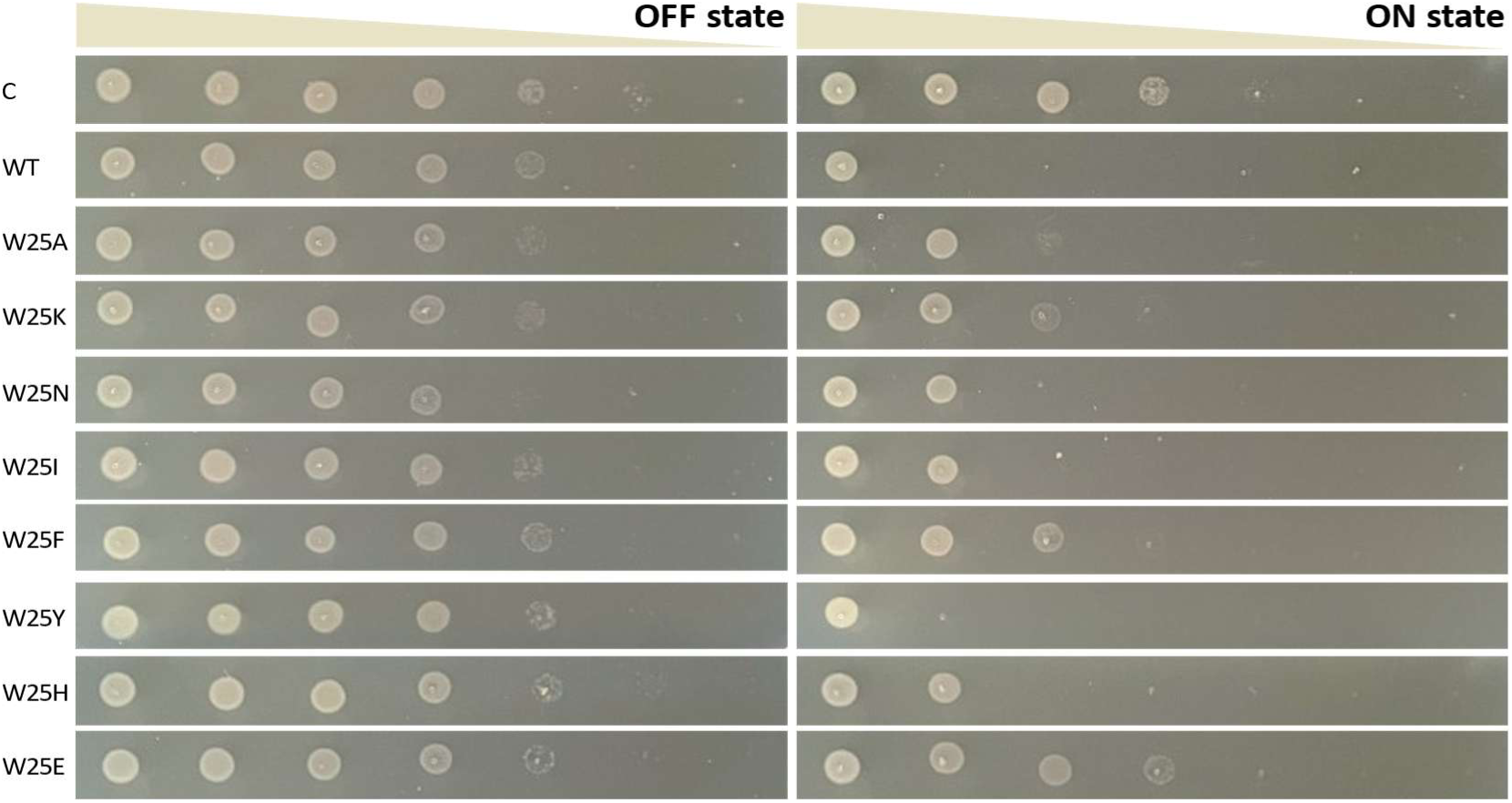
Spot test of *E. coli* EPI400 cells encoding wild-type *Vc*ParE2 (WT) and various mutants having tryptophane 25 substituted with alanine, lysine, asparagine, isoleucine, phenylalanine, tyrosine, histidine or glutamic acid, respectively. Growth of the *Vc*ParE2 variants - introduced in the P_BAD_-RS_theo_ multi-layer control system - is compared without induction (OFF state, absence of arabinose and theophylline) and upon induction (ON state, in presence of arabinose and theophylline). As a control, cells carrying the empty pET22b vector were included in the screen.

Trp25 is located in a short loop between helices α1 and α2 that form a structural hallmark of ParE’s [38]. It is nevertheless conserved only in the more closely related ParE family members, including *Mycobacterium tuberculosis* ParE1 and *Caulobacter crescentus* ParE1 (sequence identities 42.9% and 37.5% respectively), but not in the more distantly related family members *Pseudoalteromonas rubra* PrpT, *Mesorhizobium opportunistum* ParE2 and ParE3, *Shewanella oneidensis* ParE_SO_, *Escherichia coli* O157 ParE2, *Pseudomonas aeruginosa* ParE1 and *M. tuberculosis* ParE2 (sequence identities 28.3%, 20.2%, 16.2%, 16.5%, 15.2%, 14.0% and 12.1% respectively). In the latter, the loop between both helices is one residue shorter and differs in conformation, making a one-to-one identification of the residue corresponding to *Vc*ParE2 Trp25 difficult. Based on the residues present at the corresponding position in these ParE proteins, the side chain requirement at *Vc*ParE2 position 25 was therefore probed via mutation of Trp25 to Lys, Asn, Ile, Phe, Tyr, His and Glu. Substituting tryptophane at position 25 to any of these residues significantly diminishes, but does not fully eliminate, the toxic effect of *Vc*ParE2, to a similar level as seen for *Vc*ParE2^W25A^. Only the tyrosine substitution of Trp25 shows full toxicity comparable to wild-type *Vc*ParE2 (Figure 3). Thus, despite lack of conservation, Trp25 is a key residue in *Vc*ParE2.

### The C-Terminus of *Vc*ParE2 is not Involved in Toxicity

An earlier report suggested that the C-terminal flexible tail of *Caulobacter crescentus* ParE1 (*Cc*ParE1) is critical for its activity, as pMT630 and pMT552 plasmids encoding truncations of this region (*Cc*ParE1^1-92^, *Cc*ParE1^1-86^ and *Cc*ParE1^1-79^), unlike the wild-type, could be transformed into *E. coli* and *C. crescentus ΔparDE1* cells [39]. Equivalent truncates were designed for *Vc*ParE2 (VcParE2^1-93^, *Vc*ParE2^1-88^ and *Vc*ParE2^1-81^ respectively, Supplementary Table S1) based on a PROMALS 3D structure-based multiple sequence alignment with *Cc*ParE1 [40]. Contrary to the findings by Fiebig and co-workers, cells harboring the truncated *Vc*ParE2^1-88^ and *Vc*ParE2^1-93^ do not exhibit growth. Viability in truncate-carrying cells is only observed in the Trp25Ala context (VcParE2^1-88, W25A^ and *Vc*ParE2^1-93, W25A^) or by additional deletion of a significant part of the β4 strand (VcParE2^1-81^) (Supplementary Figure S2).

To further probe whether the N- or C-termini of *Vc*ParE2 are involved in possible interactions with Gyrase, two versions of a bulky SUMO-tag (full length and the folded domain consisting of residues T22-E94; Supplementary Table S1) were added to both termini. Cells harboring these constructs are not viable (Supplementary Figure S2). Additionally, in contrast to the successful co-expression of *Vc*ParE2 with *Vc*ParD2 that is achieved when *Vc*ParE2 carries an N- or C terminal His-tag (and allows for purification of *Vc*ParE2), cells also cannot grow when an N- or C- terminally SUMO-tagged or Twin-Strep-tagged version *Vc*ParE2 is co-expressed with *Vc*ParD2. The SUMO- and Twin-Strep-tags thus do not seem to block its interaction with Gyrase, but do interfere with the formation of the *Vc*ParE2-VcParD2 complex.

### Important Players in *Vc*ParE2 Toxicity are Located at the α1-α2 Hairpin

Protein activity often is the result of an interplay among different residues, rather than just one amino acid. To identify potential other residues associated with *Vc*ParE2 toxicity, the preliminary screen was expanded with multiple alanine mutants combining the Trp25Ala mutation with additional alanine substitutions of surrounding residues. Growth assessment was performed on these *Vc*ParE2-variants introduced in the Ptac-RS_theo_ multi-layer control system [36]. The latter system was here chosen over the P_BAD_-RS_theo_ system to allow more subtle differences in toxicity but has the disadvantage that no wild-type parE2 can be cloned therein. These experiments reveal that an additional alanine-replacement of residues Phe20, Tyr33, His69 and His88, on top of the Trp25Ala mutation, have the largest effect on colony growth (Figure 4). Tyr33 and His69 are partially buried, and therefore may also have a structural role, while Phe20 and His88 are exposed residues, with His88 showing high conservation. Spot tests show how every additional mutation causes a gradual decrease in in vivo toxicity upon induction: *Vc*ParE2^F20A, W25A^ is less toxic than *Vc*ParE2^W25A^ (Figure 5A). The *Vc*ParE2^F20A, H88A^ double mutant reduces toxicity to a level comparable to what is observed for *Vc*ParE2^W25A^ (Figure 5A).

**Figure 4:**
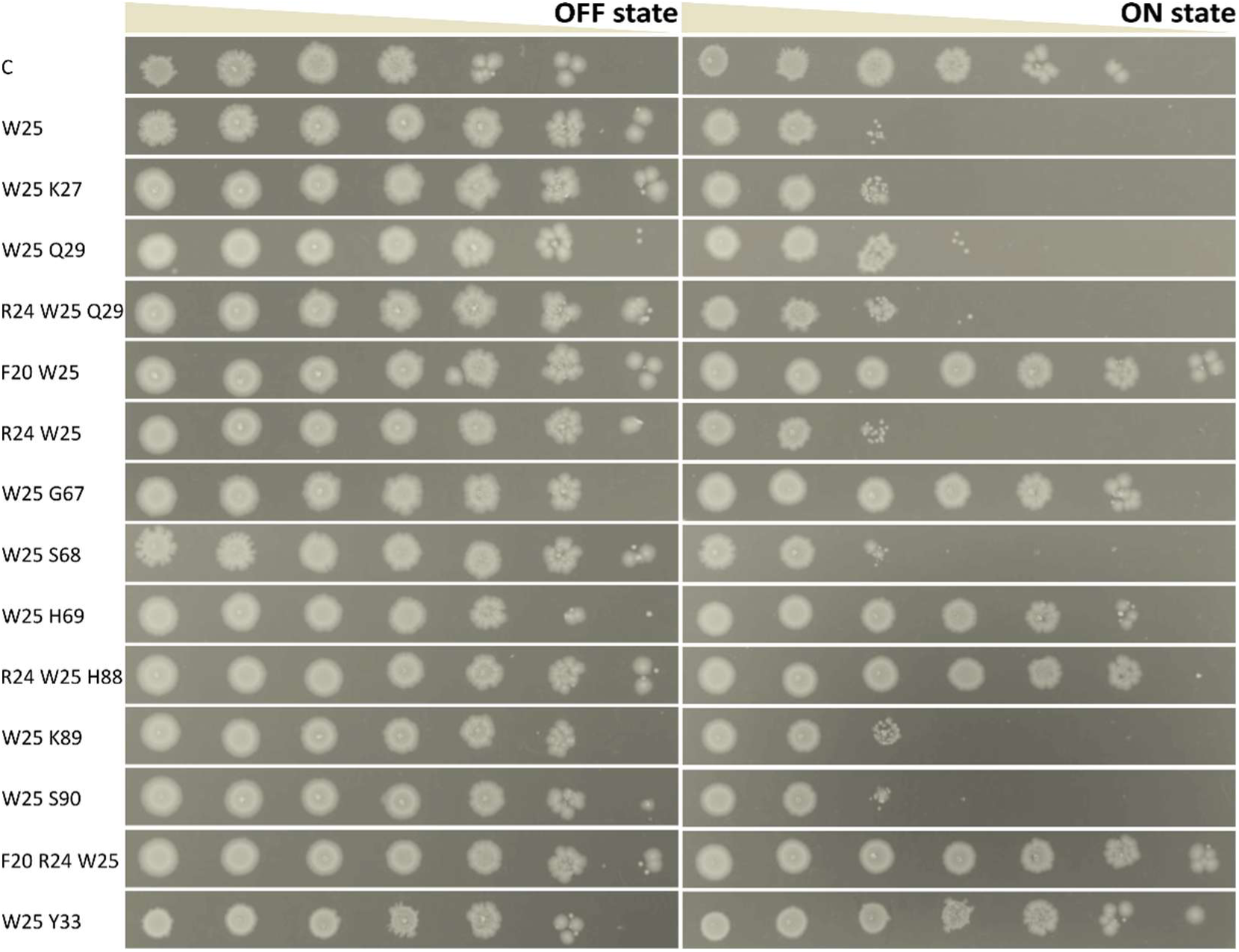
Spot test of *E. coli* EPI400 cells encoding multiple alanine mutants of *Vc*ParE2. Growth of the *Vc*ParE2 variants - introduced in the P*tac*-RS_theo_ multi-layer control system - is compared without induction (OFF state, absence of IPTG and theophylline) and upon induction (ON state, in presence of IPTG and theophylline). As a control, cells carrying the empty pET22b vector (C) and the Trp25Ala single alanine mutation (W25) were included in the screen.

**Figure 5:**
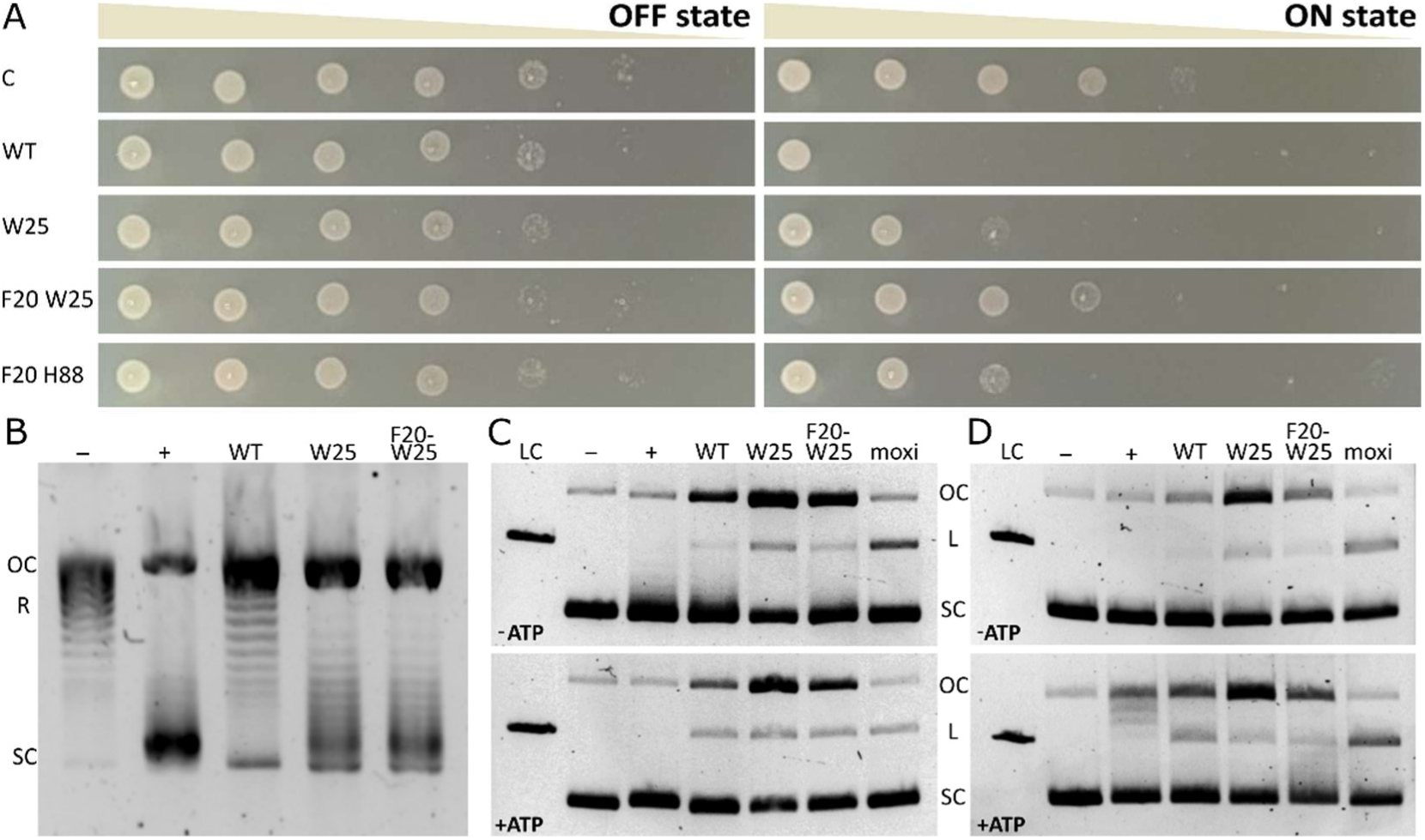
Effect of *Vc*ParE2 mutations on its toxic activity. (**A**) Spot tests assessing the growth of *E. coli* EPI400 cells expressing various *Vc*ParE2 variants under the P_BAD_-RS_theo_ multi-layer control system. Growth was compared under ON conditions (toxin expression induced by arabinose and theophylline) and OFF conditions (no induction). Tested variants include wild-type *Vc*ParE2 (WT), *Vc*ParE2^W25A^ (W25), *Vc*ParE2^F20A, W25A^ (F20 W25) and *Vc*ParE2^F20A, H88A^ (F20 H88). As a non-toxic control, *E. coli* EPI400 cells carrying an empty pET22b vector (**C**) were included. (**B**) Gyrase supercoiling assay evaluating the ability of wild-type *Vc*ParE2 (WT), single mutant *Vc*ParE2^W25A^ (W25) and double mutant *Vc*ParE2^F20A, W25A^ (F20 W25) to introduce negative supercoils into relaxed pBR322 plasmid. DNA topoisomers are labelled as OC (open circular), R (relaxed) and SC (supercoiled). (**C**) Gyrase cleavage assays assessing the stabilization of the Gyrase-DNA cleavage complex in presence of wild-type *Vc*ParE2 (WT), single mutant *Vc*ParE2^W25A^ (W25) and double mutant *Vc*ParE2^F20A, W25A^ (F20 W25). Reactions were carried out in absence and in presence of ATP. Moxifloxacin was used as control. DNA topoisomers are labelled as OC (open circular), L (linear) and SC (supercoiled). (**D**) Same as (C) but using Topoisomerase IV instead of Gyrase. The left lane (LC) represents linearized pBR322. Negative control lanes (-) contain only DNA (relaxed pBR322 for the supercoiling assays, negatively supercoiled pBR322 for the cleavage assays), positive control lanes (+) contain both DNA and Gyrase (B,C) or topoisomerase IV (D).

Wild-type *Vc*ParE2, *Vc*ParE2^W25A^ and *Vc*ParE2^F20A, W25A^ were co-expressed with *Vc*ParD2 in *E. coli* and purified using a previously established protocol [32]. All three proteins elute as single peaks during analytical gel filtration, with therefrom derived estimated molecular weights of 8.1 kDa, 9.5 kDa and 9.9 kDa respectively, indicating that they all behave as a monomer in solution (Supplementary Figure S3A). Circular dichroism data confirm that all variants are well-folded, with the overall secondary structure of the toxin conserved, despite the introduction of one or two alanine substitutions (Supplementary Figure S3B). Therefore, the decrease in activity is not attributable to a loss of structure caused by the mutations.

Next, we assessed the effects of the *Vc*ParE2 mutations on the ability of Gyrase to negatively supercoil pBR322. While wild-type *Vc*ParE2 inhibits Gyrase supercoiling, both mutants exhibit significantly reduced supercoiling activity (Figure 5B). Nevertheless, all variants stabilize the cleavage complex of both Gyrase and Topoisomerase IV in the presence of ATP. Stabilization is also observed in the absence of ATP, albeit to a lesser extent (Figure 5C-D, Supplementary Figure S4). This suggests that the Trp25A (and Phe20A) mutation does not fully abolish supercoiling activity, an observation that resembles the effect that was earlier observed for Microcin B17 [41]. It thus is possible that the mutations affect the efficiency of cleavage and relegation, slowing down the supercoiling process rather than stopping it entirely.

The full C-Terminus of *Vc*ParD2 is Required to Inhibit *Vc*ParE2 in vivo and in vitro *Vc*ParD2 consists of an N-terminal DNA-binding domain that adopts a ribbon-helix-helix fold, while its C-terminal domain is unstructured in absence of *Vc*ParE2, but undergoes folding upon binding to *Vc*ParE2. The interaction between *Vc*ParD2 and *Vc*ParE2 results in a 6:2 complex with an apparent dissociation constant (k_D_) of 45 nM [32]. Biolayer interferometry confirmed the binding of *Vc*ParE2 to the C-terminal domain of *Vc*ParD2 (Supplementary Figure S5A). However, precise affinity determination was not possible due to the release of the complex from the biosensor during the association step. The reason thereof remains unclear. Binding between *Vc*ParD2 and *Vc*ParE2 appears to be initiated by the N-terminal helix of the C-terminal domain (ParD2^E41-S60^), as this fragment binds *Vc*ParE2 with a K_D_ of 778 nM (Supplementary Figure S5A; Figure 6A). In contrast, no binding was observed between *Vc*ParE2 and the C-terminal region of the *Vc*ParD2 C-terminal domain (ParD2^G61-R80^) (Supplementary Figure S5C-D).

**Figure 6:**
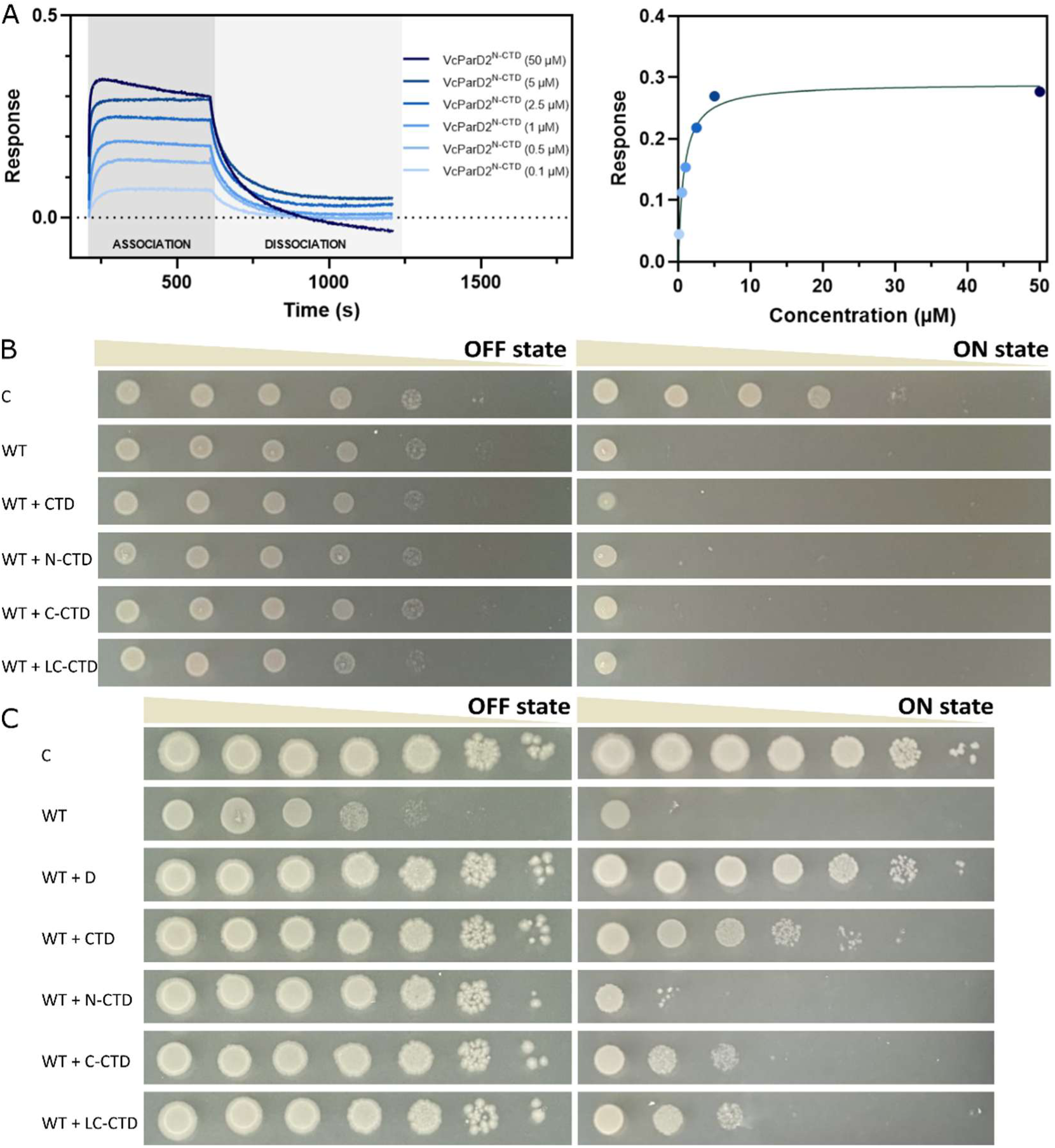
Effect of *Vc*ParD2 and its fragments on *Vc*ParE2 toxicity. (**A**) BLI curves depicting the interaction between immobilized *Vc*ParE2 and the N-terminal helix of the *Vc*ParD2 C-terminal domain (VcParD2^E41-S60^). Steady-state analysis of this data set indicates a moderate binding affinity (778 nM) between *Vc*ParE2 and the N-terminal helix of the *Vc*ParD2 C-terminus. (**B-C**) Spot test of *E. coli* EPI400 cells expressing wild-type *Vc*ParE2 (WT) under control of the P_BAD_-RS_theo_ multi-layer control system, either *in cis* (B) or *in trans* (**C**) with various *Vc*ParD2 subfragments: *Vc*ParD2^E41-R80^ (CTD), *Vc*ParD2^E41-S60^ (N-CTD), *Vc*ParD2^D66-R80^ (C-CTD), *Vc*ParD2^G61-R80^ (LC-CTD). In the in cis configuration, antitoxin subfragments are co-expressed with the toxin under the same P_BAD_ promoter within the bicistronic operon. In contrast, the *in trans system* places antitoxin fragments under the control of a strong tac promoter. For both conditions, cell growth was compared either switching toxin production OFF (absence of arabinose and theophylline) or ON (in presence of arabinose and theophylline). As a control, cells harboring the empty pET22b vector (C) were included in the assay.

Spot tests indicate that in cis co-expression of C-terminal *Vc*ParD2 fragments with *Vc*ParE2, mimicking the antitoxin:toxin ratio obtained in the bicistronic *par*DE operon, is insufficient to neutralize toxicity *in vivo*. When, on the other hand, the same fragments are over-expressed *in trans*, only full-length *Vc*ParD2 fully inhibits *Vc*ParE2. The C-terminal domain of the antitoxin *Vc*ParD2 partially prevents *Vc*ParE2 toxicity, whereas smaller subfragments exhibit significantly less or even no protection (Figure 6B,C). Altogether, the N-terminal region of the C-terminus contributes to antitoxin binding, but the entire C-terminus is required for inhibition of *Vc*ParE2 toxicity, along with the folded N-terminal domain to secure *Vc*ParE2 within the TA complex.

## DISCUSSION

VcParE2 is a member of RelE/ParE toxin superfamily. Although RelE and ParE toxins share little sequence identity and have distinct biochemical activities (ParE acts as a Gyrase poison, while RelE functions as a ribosome-dependent mRNase), they adopt a very similar structure consisting of two N-terminal α-helices followed by four anti-parallel β-strands. Nevertheless, two distinct structural features distinguish the ParE and RelE family members. First of all, the N-terminal helices of ParE’s are significantly longer than those of RelE’s. secondly, RelE’s contain an additional α-helix at their C-terminus which seems to be absent in ParE structures: the C-terminus of ParE is shorter and, in contrast to RelE, not predicted to adopt a helix structure (Figure 7A-B).

**Figure 7:**
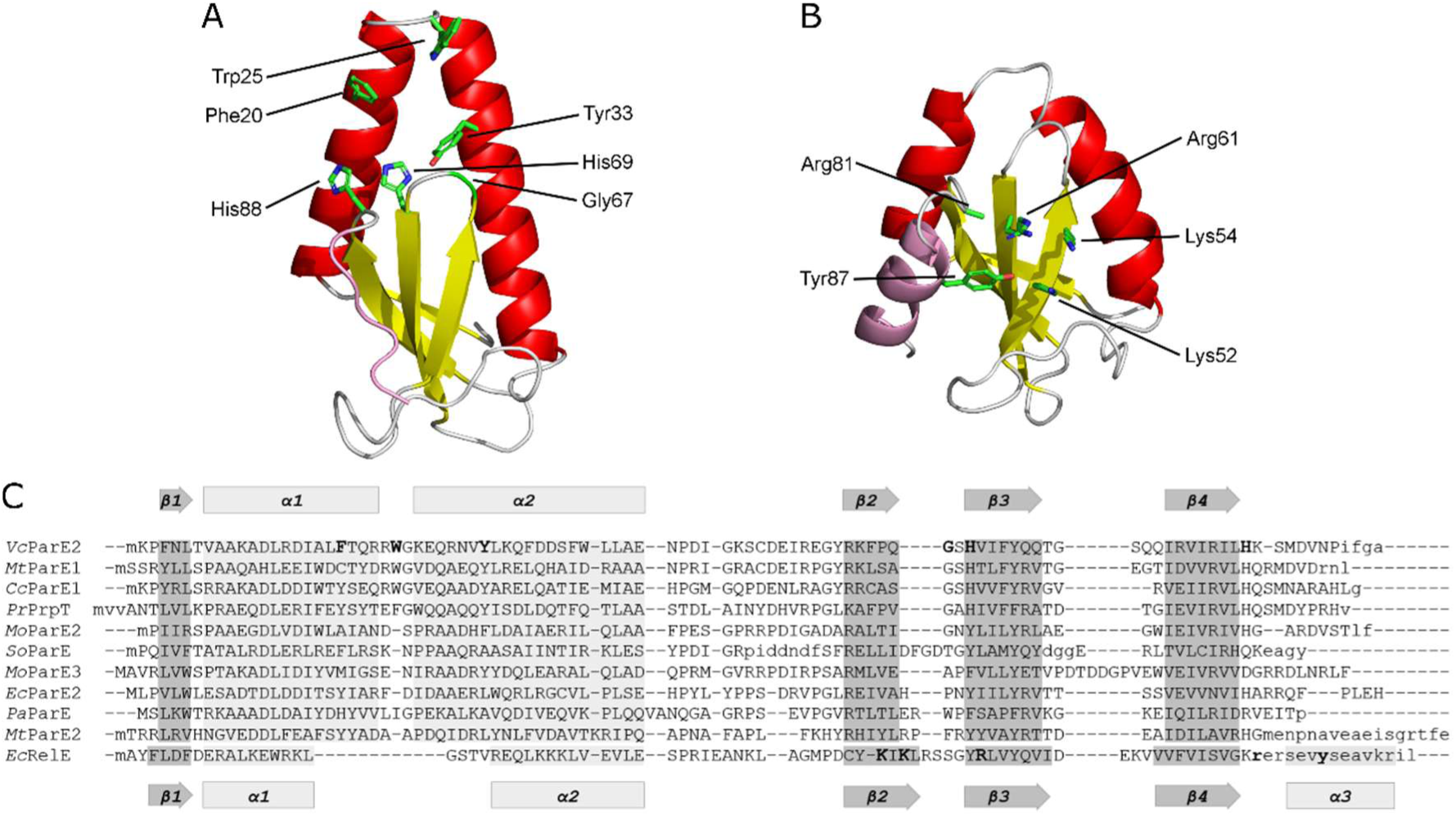
Structural comparison between the ParE and RelE families. (**A**) Cartoon representation of *Vc*ParE2 (PDB ID: 7R5A). β-sheets are colored yellow and α-helices in red. Residues important for toxicity are indicated in green stick representation and labelled. The C-terminus is indicated in pink. (**B**) Similar cartoon representation of *Ec*RelE (PDB ID: 4V7J). The active site residues are located at the β-sheet and are distinct from the key residues essential for ParE toxicity. The C-terminal α-helix that is essential for RelE activity but absent in ParE is shown in pink. (**C**) PROMALS3D structure-based alignment of ParE toxins and EcRelE. Structures used to guide the alignment are extracted from following PDB files: *Vc*ParE2 (PDB ID: 7R5A); *Mt*ParE1 (PDB ID: 8C24); *Cc*ParE1 (PDB ID: 3KXE); *Pr*PrpT (PDB ID: 7YCS); *Mo*ParE2 (lPDB ID: 6X0A); *So*ParE (PDB ID: 7ETR); *Mo*ParE3 (PDB ID: 5CEG); *Ec*ParE2 (PDB ID: 5CW7); *Pa*ParE (PDB ID: 6XRW); *Mt*ParE2 (PDB ID: 8C26); EcRelE (PDB ID: 4V7J). Residues shown in panels A and B are indicated in bold. Residues that are not visible in the crystal structures are indicated in lowercase.

It now has been demonstrated that these structural features are linked with their respective activities. Previous studies established that the C-terminal helix of RelE and related RNase toxins, next to catalytic residues on the β-sheet, is essential for ribosome-dependent mRNA cleavage [42]. Data presented here demonstrates that the activity of *Vc*ParE2 is crucially dependent on amino acids located at the tip of the α-hairpin formed by the two N-terminal α-helices. Thus, both activities (RNase and Gyrase inhibition) are located at distinct regions of the RelE/ParE fold and allow for a mechanism by which both toxin families may have evolved from a common ancestor, without having to pass over an inactive intermediate. Likely, this ancestor was a RelE-like ribonuclease, given that the RelE family (which also includes a.o. YoeB, HigB, MqsR, YafQ, YafO, Yhav) is larger and more diverse than the ParE family. This hypothesis is further supported by a recent phylogenetic analysis of the RelE/ParE superfamily, where it was discovered that the ParE family includes a subcluster of mRNA-cleaving toxins [43].

Although earlier studies suggest that the C-terminus of ParE is important for its toxicity [39,44,30], we showed that truncation of the C-terminal disordered segment of *Vc*ParE2 does not affect its toxicity, nor does the addition of a bulky protein or tag. This supports the recent findings of Bourne and colleagues, who demonstrated that adding the C-terminus of toxic ParE toxins (VcParE2 and *Cc*ParE1) to a non-toxic one (*Pa*ParE1) did not induce toxicity [45]. The most crucial residue for *V. cholerae* ParE2 toxicity was identified to be Trp25, as alanine substitution of Trp25 implies a decrease in toxicity. Nevertheless, this residue is not well conserved within the ParE family (Figure 7C). While present in some ParE’s that are quite closely related to *V. cholerae* ParE2, *Mycobacterium tuberculosis* ParE1 (42.9% sequence identity) and *Caulobacter crescentus* ParE1 (37.5% sequence identity), most ParE’s carry residues different from tryptophane at equivalent positions (Lys in *S. oneidensis*, Asn in *M. opportunistum*, Ile in *P. aeruginosa*, Phe in *E. coli* 0157 ParE2). Substitution of *Vc*ParE2(Trp25) in into these residues creates variants with diminished toxicity.

A comparable situation is observed for the otherwise unrelated Gyrase poison CcdB encoded by the F-plasmid (CcdB^F^), where the tryptophan residue at position 99 stacks with Arg462 in the Gyrase A14 fragment [46]. This forms a key interaction in the binding mechanism between CcdB and Gyrase. Substitution of Trp99 in CcdB^F^ results in a complete loss of toxicity. Still, Trp99 is not conserved across CcdB homologues. *Vibrio fischeri* CcdB (CcdB^vfi^), for instance, hosts Thr at this position. Despite using the same binding interface, CcdB^vfi^ employs alternative interactions to bind Gyrase, where Gyrase interacts with Asp99^vfi^ instead of Thr103^vfi^ - the residue positioned equivalently to Trp99^F^ [47].

Different ParD-ParE complexes are similar in how the C-terminal domain of ParD interacts with ParE, but heavily differ in the global architecture and stoichiometry of the complex. The C-terminal halve of *Vc*ParD2 (VcParD2^41-80^), which offers partial protection against *Vc*ParE2, wraps around the toxin and interacts with the α-hairpin, but does not cover Trp25. Likely, this *Vc*ParD2^41-80^ segment prevents interaction between Gyrase and *Vc*ParE2 via steric hindrance. Nevertheless, *Vc*ParD2^41-80^ provides only partial protection against *Vc*ParE2. Dissociation of the *Vc*ParE2-VcParD2^41-80^ complex at the presumed very low concentrations in the cell is still sufficient to allow competition with Gyrase for *Vc*ParE2 binding. Full protection against *Vc*ParE2 is only obtained in presence of full-length *Vc*ParD2, capturing the toxin in a heterooctameric complex (PDB ID: 7R5A, [48]). In the heterooctameric assembly, two *Vc*ParE2 monomers contact each other, thereby shielding Trp25 from the solvent and at the same time further stabilizing the *Vc*ParE2-VcParD2 complex. Correct formation of this architecture is likely hampered by adding a bulky SUMO tag to either the N- or C-terminus of *Vc*ParE2, as the latter prevents it from being protected by *Vc*ParD2.

The quaternary arrangement of other ParD-ParE complexes for which structures are available deviates from the 6:2 complex observed for *Vc*ParDE2 [28, 38, 48–54]. Although the exact arrangement varies from case to case, most ParDE-complexes adopt a heterotetrameric form in which two ParE toxins contact each other. When the *Vc*ParD2-to-VcParE2 ratio drops, such heterotetramers are also formed in case of VcparDE2, yet the exact architecture here remains unknown. Similar to what is seen for *Vc*ParE2, residues equivalent to *Vc*ParE2(Trp25) in *Cc*ParE1, *Ec*ParE2, *Mo*ParE2, *Mo*ParE3, *Mt*ParE1, *Mt*ParE2, *Pa*ParE, *Pr*PrpT, *So*ParE are exposed to the solvent in the free toxin state. However, in the different complexes with their cognate ParD, they become (partially) buried, independent of the exact architecture of the complex (Figure 8). Thus, the “strategy” to protect the cell against accidental ParE activation seems to be a combination of burying the key interaction residues of ParE combined with the use of a higher order ParD-ParE complex to obtain a highly stable complex. Such a strategy has been observed for other, unrelated toxin-antitoxin systems as well, an example being *faRel2/aTfaRel2* [55].

**Figure 8:**
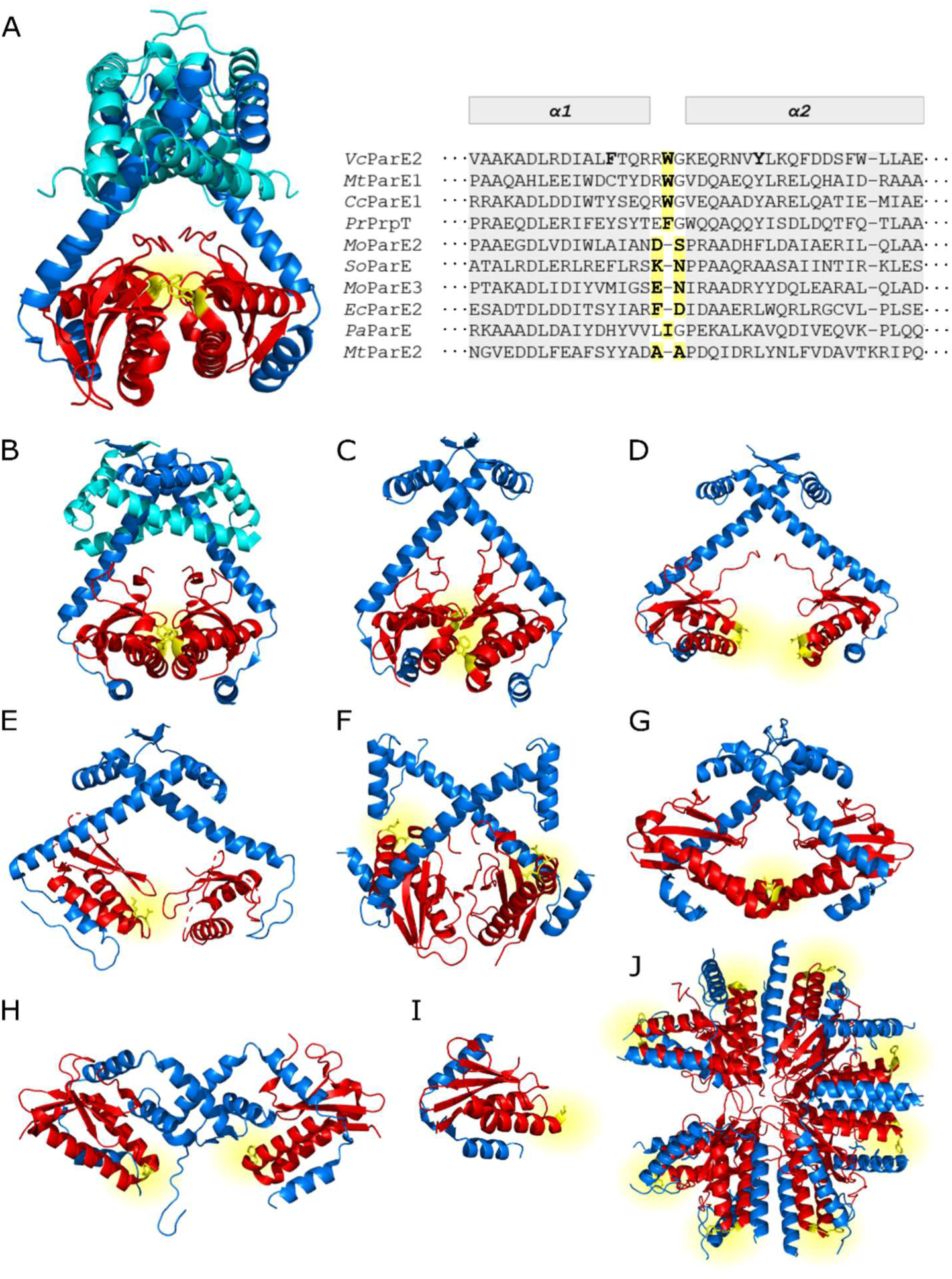
Residue Trp25 of *Vc*ParE2 and its equivalent positions in homologous ParE proteins, marked in yellow in the protein sequences (right upper corner) and on the currently available 3D structures of ParDE TA-complexes (A-J). Antitoxins are colored blue and cyan, toxins red. (**A**) *Vibrio cholerae* ParDE2; (**B**) *Mycobacterium tuberculosis* ParDE1; (**C**) *Caulobacter crescentus* ParDE1; (**D**) *Mhezorizobium opportunistum* ParDE2; (**E**) *Shewanella oneidensis* ParE_SO_- CopA_SO_; (**F**) *Mesorhizobium opportunistum* ParDE3; (**G**) *Pseudomonas aeruginosa* ParDE1; (**H**) *Pseudoalteromonas rubra* PrpAT; (**I**) *Mycobacterium tuberculosis* ParDE2; (**J**) *Escherichia coli* PaaA2-ParE2.

## ACKNOWLEDGEMENTS

We sincerely thank Prof. dr. Anthony Maxwell (John Innes Centre, United Kingdom) for his insights in the supercoiling and cleavage assays, and Prof. dr. em. Henri De Greve for his advice on the cloning and spot tests. We thank Nisrine El Mesbahi for her practical contributions to the single alanine scan.

## AUTHOR CONTRIBUTIONS

Yana Girardin: Conceptualization, Funding acquisition, Formal analysis, Investigation, Methodology, Validation, Visualization, writing - original draft, writing - review and editing Remy Loris: Conceptualization, Funding acquisition, Methodology, Resources, writing - original draft, writing - review and editing

## CONFLICT OF INTEREST

N.A.

## FUNDING

This work was supported by a personal PhD mandate to Y.G. from FWO [1164622N] and to grant OZR4292 from OZR-VUB to R.L.

## Notes

### Competing Interest Statement

The authors have declared no competing interest.

