## Supplementary figures for "Inhibition of Gyrase via a conserved α-hairpin in *Vibrio cholerae* ParE2 and neutralization by ParD2"

**Supplementary Table S1:** List of *Vc*ParE2 synthetic fragments and synthetic fragments for co-expression of *Vc*ParD2 with SUMO-coupled *Vc*ParE2. All *Vc*ParE2 single alanine mutants are variants of the wild-type *Vc*ParE2 fragment listed in this table, with the codon of the targeted residue replaced by GCC. Overlapping regions with the pET28b vector are written in lowercase, while the *Vc*ParD2 and *Vc*ParE2 sequences are highlighted in blue and red uppercase, respectively. The full-length SUMO (SUMO$^{FL}$) and SUMO folded domain (SUMO$^{T22\text{-}E94}$) sequences are shown in grey uppercase. The start codon is indicated in bold and NcoI and HindIII restriction sites are underlined. Like wild-type *VcparDE2*, the *VcparD2* coding sequence shows a 4-bp overlap with the subsequent gene (brown). (*continued on next page*)

| Name | Sequence insert (5′ → 3′) |
|---|---|
| *Vc*ParE2 WT | gtttaactttaagaaggagatatacc**atg**GGCAAACCATTTAATCTTACCGTCGCCGCCAAAGCCGATTTACGTGATATTGCTT<br>TATTCACTCAACGACGCTGGGGAAAAGAGCAGCGAAATGTTTATTTAAAGCAATTCGATGATTCCTTTTGGCTTTTAG<br>CGGAAAATCCCGACATTGGTAAATCATGCGATGAAATCCGAGAGGGATACAGAAAATTTCCCCAAGGGAGTCACGT<br>CATCTTTTATCAGCAAACCGGCAGCCAACAAATCAGGGTGATCCGAATTCTTCATAAGAGCATGGATGTGAACCCAA<br>TATTCGGCGCACACCATCACCATCACCATTGAaagcttgcggccgcactcgagcac |
| SUMO$^{FL}$-*Vc*ParE2 | gtttaactttaagaaggagatatacc**atg**GGCAGCGATAGCGAGGTGAACCAGGAAGCGAAGCCGGAAGTGAAACCGGAA<br>GTTAAGCCGGAAACCCACATCAACCTGAAAGTTAGCGACGGTAGCAGCGAGATCTTCTTTAAGATTAAGAAAACCA<br>CCCCGCTGCGTCGTCTGATGGAAGCGTTCGCGAAACGTCAGGGCAAGGAGATGGACAGCCTGCGTTTTCTGTACGA<br>TGGCATCCGTATTCAAGCGGACCAGACCCCGGAAGACCTGGATATGGAGGACAACGATATTATTGAAGCGCATCGTG<br>AACAAATCGGCGGCGGCAGCGGCATGAAACCATTTAATCTTACCGTCGCCGCCAAAGCCGATTTACGTGATATTGCT<br>TTATTCACTCAACGACGCTGGGGAAAAGAGCAGCGAAATGTTTATTTAAAGCAATTCGATGATTCCTTTTGGCTTTTA<br>GCGGAAAATCCCGACATTGGTAAATCATGCGATGAAATCCGAGAGGGATACAGAAAATTTCCCCAAGGGAGTCACG<br>TCATCTTTTATCAGCAAACCGGCAGCCAACAAATCAGGGTGATCCGAATTCTTCATAAGAGCATGGATGTGAACCCA<br>ATATTCGGCGCACACCATCACCATCACCATTGAaagcttgcggccgcactcgagcac |
| *Vc*ParE2-SUMO$^{FL}$ | gtttaactttaagaaggagatatacc**atg**GGCAAACCATTTAATCTTACCGTCGCCGCCAAAGCCGATTTACGTGATATTGCTT<br>TATTCACTCAACGACGCTGGGGAAAAGAGCAGCGAAATGTTTATTTAAAGCAATTCGATGATTCCTTTTGGCTTTTAG<br>CGGAAAATCCCGACATTGGTAAATCATGCGATGAAATCCGAGAGGGATACAGAAAATTTCCCCAAGGGAGTCACGT<br>CATCTTTTATCAGCAAACCGGCAGCCAACAAATCAGGGTGATCCGAATTCTTCATAAGAGCATGGATGTGAACCCAA<br>TATTCGGCGCAATGAGCGATAGCGAGGTGAACCAGGAAGCGAAGCCGGAAGTGAAACCGGAAGTTAAGCCGGAA<br>ACCCACATCAACCTGAAAGTTAGCGACGGTAGCAGCGAGATCTTCTTTAAGATTAAGAAAACCACCCCGCTGCGTCG<br>TCTGATGGAAGCGTTCGCGAAACGTCAGGGCAAGGAGATGGACAGCCTGCGTTTTCTGTACGATGGCATCCGTATTC<br>AAGCGGACCAGACCCCGGAAGACCTGGATATGGAGGACAACGATATTATTGAAGCGCATCGTGAACAAATCGGCG<br>GCGGCAGCGGCCACCACCACCACCACTGAaagcttgcggccgcactcgagcac |
| SUMO$^{T22\text{-}E94}$-*Vc*ParE2 | gtttaactttaagaaggagatatacc**atg**GGCACCCACATCAACCTGAAAGTTAGCGACGGTAGCAGCGAGATCTTCTTTAAG<br>ATTAAGAAAACCACCCCGCTGCGTCGTCTGATGGAAGCGTTCGCGAAACGTCAGGGCAAGGAGATGGACAGCCTG<br>CGTTTTCTGTACGATGGCATCCGTATTCAAGCGGACCAGACCCCGGAAGACCTGGATATGGAGGACAACGATATTAT<br>TGAAGCGCATCGTGAAATGAAACCATTTAATCTTACCGTCGCCGCCAAAGCCGATTTACGTGATATTGCTTTATTCACT<br>CAACGACGCTGGGGAAAAGAGCAGCGAAATGTTTATTTAAAGCAATTCGATGATTCCTTTTGGCTTTTAGCGGAAA<br>ATCCCGACATTGGTAAATCATGCGATGAAATCCGAGAGGGATACAGAAAATTTCCCCAAGGGAGTCACGTCATCTTT<br>TATCAGCAAACCGGCAGCCAACAAATCAGGGTGATCCGAATTCTTCATAAGAGCATGGATGTGAACCCAATATTCGG<br>CGCACACCATCACCATCACCATTGAaagcttgcggccgcactcgagcac |
| *Vc*ParE2-SUMO$^{T22\text{-}E94}$ | gtttaactttaagaaggagatatacc**atg**GGCAAACCATTTAATCTTACCGTCGCCGCCAAAGCCGATTTACGTGATATTGCTT<br>TATTCACTCAACGACGCTGGGGAAAAGAGCAGCGAAATGTTTATTTAAAGCAATTCGATGATTCCTTTTGGCTTTTAG<br>CGGAAAATCCCGACATTGGTAAATCATGCGATGAAATCCGAGAGGGATACAGAAAATTTCCCCAAGGGAGTCACGT<br>CATCTTTTATCAGCAAACCGGCAGCCAACAAATCAGGGTGATCCGAATTCTTCATAAGAGCATGGATGTGAACCCAA<br>TATTCGGCGCAATGACCCACATCAACCTGAAAGTTAGCGACGGTAGCAGCGAGATCTTCTTTAAGATTAAGAAAACC<br>ACCCCGCTGCGTCGTCTGATGGAAGCGTTCGCGAAACGTCAGGGCAAGGAGATGGACAGCCTGCGTTTTCTGTACG<br>ATGGCATCCGTATTCAAGCGGACCAGACCCCGGAAGACCTGGATATGGAGGACAACGATATTATTGAAGCGCATCGT<br>GAACACCACCACCACCACCACTGAaagcttgcggccgcactcgag |

**Table 5.2 continued**: The Twin-Strep-tag is underlined in the *Vc*ParE2 sequence.

| Name | Sequence insert (5′ → 3′) |
|---|---|
| *Vc*ParD2-*Vc*ParE2-Strep | gtttaactttaagaaggagatatacc**atg**GCTAAAAATACAAGTATCACTCTTGGTGAACACTTCGATGGCTTTATTACAAGCC<br>AAATACAAAGTGGGCGTTACGGCTCAGCAAGTGAAGTCATTCGCTCTGCGCTACGTCTACTCGAAAACCAAGAAAC<br>CAAACTACAGTCACTCCGTCAACTACTTATTGAAGGAGAGCAAAGTGGTGACGCTGATTATGACCTTGATAGCTTCAT<br>CAATGAACTCGATAGTGAAAACATTCGATGAAACCATTTAATCTTACCGTCGCCGCCAAAGCCGATTTACGTGATATT<br>GCTTTATTCACTCAACGACGCTGGGGAAAAGAGCAGCGAAATGTTTATTTAAAGCAATTCGATGATTCCTTTTGGCTT<br>TTAGCGGAAAATCCCGACATTGGTAAATCATGCGATGAAATCCGAGAGGGATACAGAAAATTTCCCCAAGGGAGTC<br>ACGTCATCTTTTATCAGCAAACCGGCAGCCAACAAATCAGGGTGATCCGAATTCTTCATAAGAGCATGGATGTGAAC<br>CCAATATTCGGCGCATGGTCTCACCCTCAGTTCGAAAAGGGTGGTGGCTCTGGTGGTGGTTCTGGTGGTGGTTCTTG<br>GTCCCACCCCCAGTTCGAGAAGTGAaagcttgcggccgcactcgagcac |
| *Vc*ParD2-SUMO$^{FL}$-*Vc*ParE2 | gtttaactttaagaaggagatatacc**atg**GCTAAAAATACAAGTATCACTCTTGGTGAACACTTCGATGGCTTTATTACAAGCC<br>AAATACAAAGTGGGCGTTACGGCTCAGCAAGTGAAGTCATTCGCTCTGCGCTACGTCTACTCGAAAACCAAGAAAC<br>CAAACTACAGTCACTCCGTCAACTACTTATTGAAGGAGAGCAAAGTGGTGACGCTGATTATGACCTTGATAGCTTCAT<br>CAATGAACTCGATAGTGAAAACATTCGATGAGCGATAGCGAGGTGAACCAGGAAGCGAAGCCGGAAGTGAAACCG<br>GAAGTTAAGCCGGAAACCCACATCAACCTGAAAGTTAGCGACGGTAGCAGCGAGATCTTCTTTAAGATTAAGAAAA<br>CCACCCCGCTGCGTCGTCTGATGGAAGCGTTCGCGAAACGTCAGGGCAAGGAGATGGACAGCCTGCGTTTTCTGTA<br>CGATGGCATCCGTATTCAAGCGGACCAGACCCCGGAAGACCTGGATATGGAGGACAACGATATTATTGAAGCGCAT<br>CGTGAAATGAAACCATTTAATCTTACCGTCGCCGCCAAAGCCGATTTACGTGATATTGCTTTATTCACTCAACGACGCT<br>GGGGAAAAGAGCAGCGAAATGTTTATTTAAAGCAATTCGATGATTCCTTTTGGCTTTTAGCGGAAAATCCCGACATT<br>GGTAAATCATGCGATGAAATCCGAGAGGGATACAGAAAATTTCCCCAAGGGAGTCACGTCATCTTTTATCAGCAAAC<br>CGGCAGCCAACAAATCAGGGTGATCCGAATTCTTCATAAGAGCATGGATGTGAACCCAATATTCGGCGCACATCATC<br>ATCATCATCACTGAaagcttgcggccgcactcgagcac |
| *Vc*ParD2-*Vc*ParE2-SUMO$^{FL}$ | gtttaactttaagaaggagatatacc**atg**GCTAAAAATACAAGTATCACTCTTGGTGAACACTTCGATGGCTTTATTACAAGCC<br>AAATACAAAGTGGGCGTTACGGCTCAGCAAGTGAAGTCATTCGCTCTGCGCTACGTCTACTCGAAAACCAAGAAAC<br>CAAACTACAGTCACTCCGTCAACTACTTATTGAAGGAGAGCAAAGTGGTGACGCTGATTATGACCTTGATAGCTTCAT<br>CAATGAACTCGATAGTGAAAACATTCGATGAAACCATTTAATCTTACCGTCGCCGCCAAAGCCGATTTACGTGATATT<br>GCTTTATTCACTCAACGACGCTGGGGAAAAGAGCAGCGAAATGTTTATTTAAAGCAATTCGATGATTCCTTTTGGCTT<br>TTAGCGGAAAATCCCGACATTGGTAAATCATGCGATGAAATCCGAGAGGGATACAGAAAATTTCCCCAAGGGAGTC<br>ACGTCATCTTTTATCAGCAAACCGGCAGCCAACAAATCAGGGTGATCCGAATTCTTCATAAGAGCATGGATGTGAAC<br>CCAATATTCGGCGCACATCACCATCACCATCACAGCGATAGCGAGGTGAACCAGGAAGCGAAGCCGGAAGTGAAA<br>CCGGAAGTTAAGCCGGAAACCCACATCAACCTGAAAGTTAGCGACGGTAGCAGCGAGATCTTCTTTAAGATTAAGA<br>AAACCACCCCGCTGCGTCGTCTGATGGAAGCGTTCGCGAAACGTCAGGGCAAGGAGATGGACAGCCTGCGTTTTCT<br>GTACGATGGCATCCGTATTCAAGCGGACCAGACCCCGGAAGACCTGGATATGGAGGACAACGATATTATTGAAGCG<br>CATCGTGAACAAATCGGCGGCGGCAGCGGCCATCACCATCACCATCACTGAaagcttgcggccgcactcgagcac |
| *Vc*ParD2-SUMO$^{T22\text{-}E94}$-*Vc*ParE2 | gtttaactttaagaaggagatatacc**atg**GCTAAAAATACAAGTATCACTCTTGGTGAACACTTCGATGGCTTTATTACAAGCC<br>AAATACAAAGTGGGCGTTACGGCTCAGCAAGTGAAGTCATTCGCTCTGCGCTACGTCTACTCGAAAACCAAGAAAC<br>CAAACTACAGTCACTCCGTCAACTACTTATTGAAGGAGAGCAAAGTGGTGACGCTGATTATGACCTTGATAGCTTCAT<br>CAATGAACTCGATAGTGAAAACATTCGATGACCCACATCAACCTGAAAGTTAGCGACGGTAGCAGCGAGATCTTCTT<br>TAAGATTAAGAAAACCACCCCGCTGCGTCGTCTGATGGAAGCGTTCGCGAAACGTCAGGGCAAGGAGATGGACAG<br>CCTGCGTTTTCTGTACGATGGCATCCGTATTCAAGCGGACCAGACCCCGGAAGACCTGGATATGGAGGACAACGATA<br>TTATTGAAGCGCATCGTGAAATGAAACCATTTAATCTTACCGTCGCCGCCAAAGCCGATTTACGTGATATTGCTTTATT<br>CACTCAACGACGCTGGGGAAAAGAGCAGCGAAATGTTTATTTAAAGCAATTCGATGATTCCTTTTGGCTTTTAGCGG<br>AAAATCCCGACATTGGTAAATCATGCGATGAAATCCGAGAGGGATACAGAAAATTTCCCCAAGGGAGTCACGTCATC<br>TTTTATCAGCAAACCGGCAGCCAACAAATCAGGGTGATCCGAATTCTTCATAAGAGCATGGATGTGAACCCAATATTC<br>GGCGCACATCATCATCATCATCACTGAaagcttgcggccgcactcgagcac |
| *Vc*ParD2-*Vc*ParE2-SUMO$^{T22\text{-}E94}$ | gtttaactttaagaaggagatatacc**atg**GCTAAAAATACAAGTATCACTCTTGGTGAACACTTCGATGGCTTTATTACAAGCC<br>AAATACAAAGTGGGCGTTACGGCTCAGCAAGTGAAGTCATTCGCTCTGCGCTACGTCTACTCGAAAACCAAGAAAC<br>CAAACTACAGTCACTCCGTCAACTACTTATTGAAGGAGAGCAAAGTGGTGACGCTGATTATGACCTTGATAGCTTCAT<br>CAATGAACTCGATAGTGAAAACATTCGATGAAACCATTTAATCTTACCGTCGCCGCCAAAGCCGATTTACGTGATATT<br>GCTTTATTCACTCAACGACGCTGGGGAAAAGAGCAGCGAAATGTTTATTTAAAGCAATTCGATGATTCCTTTTGGCTT<br>TTAGCGGAAAATCCCGACATTGGTAAATCATGCGATGAAATCCGAGAGGGATACAGAAAATTTCCCCAAGGGAGTC<br>ACGTCATCTTTTATCAGCAAACCGGCAGCCAACAAATCAGGGTGATCCGAATTCTTCATAAGAGCATGGATGTGAAC<br>CCAATATTCGGCGCACATCACCATCACCATCACACCCACATCAACCTGAAAGTTAGCGACGGTAGCAGCGAGATCTT<br>CTTTAAGATTAAGAAAACCACCCCGCTGCGTCGTCTGATGGAAGCGTTCGCGAAACGTCAGGGCAAGGAGATGGAC<br>AGCCTGCGTTTTCTGTACGATGGCATCCGTATTCAAGCGGACCAGACCCCGGAAGACCTGGATATGGAGGACAACG<br>ATATTATTGAAGCGCATCGTGAACATCACCATCACCATCACTGAaagcttgcggccgcactcgagcac |

**Supplementary Table S2:** List of mutations included in the single alanine scan. The second column shows the accessible surface area of the side chain of the corresponding residue. For each included *Vc*ParE2 mutant, the exact number of colonies after transformations was given, as well as observed mutations.

| ***Vc*ParE2 residue mutated to Ala** | **Total side chain ASA (A°$^2$)** | **Included in alanine scan?** | **Colony growth?** |
|---|---|---|---|
| M1 | 52.43 | | |
| K2 | 69.56 | YES | 0 |
| P3 | 26.44 | | |
| F4 | 6.64 | | |
| N5 | 65.59 | YES | 2 colonies carrying a frameshift mutation of *Vc*ParE2:<br>• Deletion of one nucleotide in the R24 codon<br>• Deletion of two nucleotides in the K36 codon |
| L6 | 21.56 | YES | 0 |
| T7 | 27.56 | YES | 0 |
| V8 | 131.42 | YES | 0 |
| A9 | 33.93 | | |
| A10 | 0.00 | | |
| K11 | 141.81 | YES | 2 colonies with a deletion of 10 nucleotides between the I72 and Q75 codons of *Vc*ParE2 |
| A12 | 44.60 | | |
| D13 | 17.15 | YES | 0 |
| L14 | 27.24 | YES | 0 |
| R15 | 140.31 | YES | 0 |
| D16 | 78.16 | YES | 0 |
| I17 | 4.99 | | |
| A18 | 9.31 | | |
| L19 | 91.81 | YES | 0 |
| F20 | 115.73 | YES | 0 |
| T21 | 10.08 | | |
| Q22 | 37.40 | YES | 0 |
| R23 | 54.93 | YES | 1 colony carrying a frameshift mutation of *Vc*ParE2:<br>• Deletion of one nucleotide in the M1 codon |
| R24 | 112.88 | YES | 1 colony carrying a frameshift mutation of *Vc*ParE2:<br>• Insertion of one nucleotide in the G52 codon |
| W25 | 132.01 | YES | Multiple colonies carrying *Vc*ParE2$^{W25A}$ without additional mutations/deletions/insertions. |
| G26 | | | |
| K27 | 138.67 | YES | 0 |
| E28 | 140.61 | YES | 0 |
| Q29 | 76.33 | YES | 0 |

| | | | |
|---|---|---|---|
| R30 | 46.77 | YES | 0 |
| N31 | 68.65 | YES | 0 |
| V32 | 80.24 | YES | 0 |
| Y33 | 39.23 | YES | 0 |
| L34 | 41.54 | YES | 1 colony carrying a frameshift mutation of *Vc*ParE2:<br>• Deletion of one nucleotide in the M1 codon |
| K35 | 128.09 | YES | 3 colonies carrying a frameshift mutation of *Vc*ParE2:<br>• Insertion of one nucleotide in the L45 codon (2 colonies)<br>• Insertion of one nucleotide between the R83 and V84 codon |
| Q36 | 90.90 | YES | 0 |
| F37 | 0.21 | | |
| D38 | 51.86 | YES | 0 |
| D39 | 69.35 | YES | 0 |
| S40 | 11.39 | YES | 0 |
| F41 | 4.05 | | |
| W42 | 119.37 | YES | 0 |
| L43 | 79.90 | YES | 0 |
| L44 | 0.00 | | |
| A45 | 1.82 | | |
| E46 | 97.34 | YES | 0 |
| N47 | 60.71 | YES | 0 |
| P48 | 3.63 | | |
| D49 | 80.10 | YES | 0 |
| I50 | 57.43 | YES | 0 |
| G51 | | | |
| K52 | 150.40 | YES | 0 |
| S53 | 42.52 | YES | 0 |
| C54 | 0.11 | | |
| D55 | 61.03 | YES | 0 |
| E56 | 50.00 | YES | 0 |
| I57 | 33.93 | YES | 0 |
| R58 | 35.52 | YES | 0 |
| E59 | 128.54 | YES | 0 |
| G60 | | | |
| Y61 | 54.38 | YES | 0 |
| R62 | 47.71 | YES | 0 |
| K63 | 25.59 | YES | 0 |
| F64 | 24.47 | YES | 0 |
| P65 | 82.23 | | |
| Q66 | 18.83 | YES | 0 |
| G67 | | | |
| S68 | 64.48 | YES | 0 |
| H69 | 12.97 | YES | 0 |
| V70 | 4.20 | | |
| I71 | 0.00 | | |

| | | | |
|---|---|---|---|
| F72 | 18.25 | YES | 0 |
| Y73 | 0.00 | | |
| Q74 | 45.98 | YES | 0 |
| Q75 | 48.94 | YES | 0 |
| T76 | 36.86 | YES | 0 |
| G77 | | | |
| S78 | 80.74 | YES | 0 |
| Q79 | 81.74 | YES | 0 |
| Q80 | 78.53 | YES | 0 |
| I81 | 0.00 | | |
| R82 | 83.70 | YES | 0 |
| V83 | 0.00 | | |
| I84 | 35.64 | YES | 0 |
| R85 | 85.60 | YES | 0 |
| I86 | 0.00 | | |
| L87 | 65.71 | YES | 0 |
| H88 | 115.37 | YES | 0 |
| K89 | 99.11 | YES | 0 |
| S90 | 58.12 | YES | 0 |
| M91 | 70.24 | YES | 0 |
| D92 | 82.94 | YES | 0 |
| V93 | 100.33 | YES | 0 |
| N94 | 28.37 | YES | 0 |
| P95 | 79.54 | | |
| I96 | | YES | 0 |
| F97 | | YES | 0 |
| G98 | | | |
| A99 | | | |

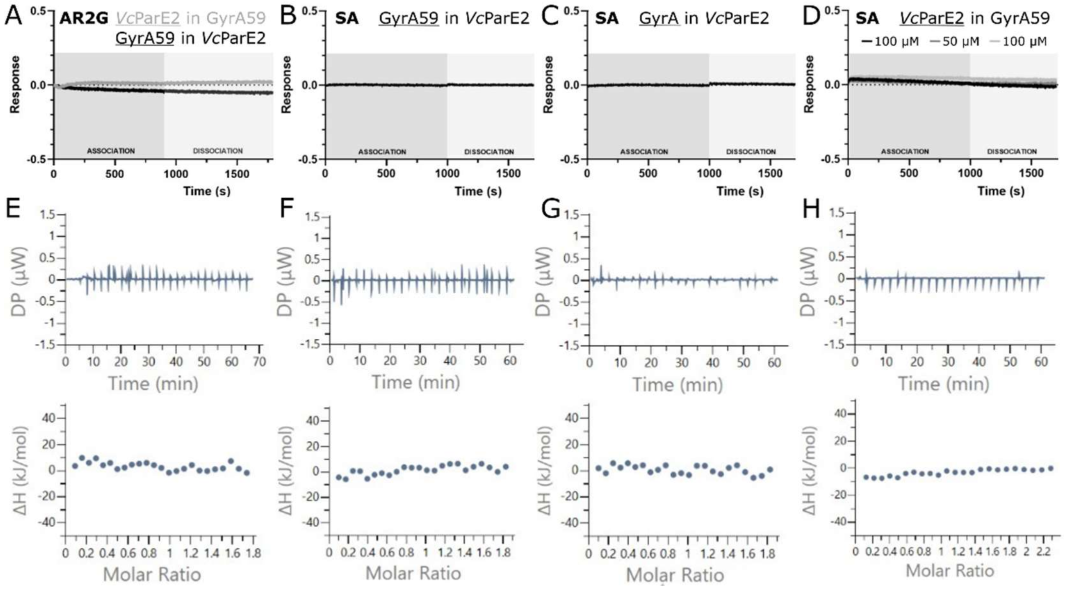


**Supplementary Figure S1**: *Vc*ParE2 does not interact with Gyrase and its subunits in absence of DNA. (**A-D**) Biolayer interferometry measurements revealed no detectable interaction between *Vc*ParE2 and either Gyrase A59 or Gyrase A, regardless of which ligand was immobilized on the biosensor (underlined). This result was consistent across different biosensor types. AR2G: proteins were immobilized via covalently coupling to the biosensor through their lysine residues. SA: Biotinylated proteins were immobilized on the streptavidin-coated biosensor surface. The association phase is represented in darker grey, while the dissociation step is shown in lighter grey. (**E-H**) Isothermal titration calorimetry. Titration of *Vc*ParE2 with Gyrase A59 (E), Gyrase A (F), Gyrase B (G) and full Gyrase (H) failed to demonstrate binding.

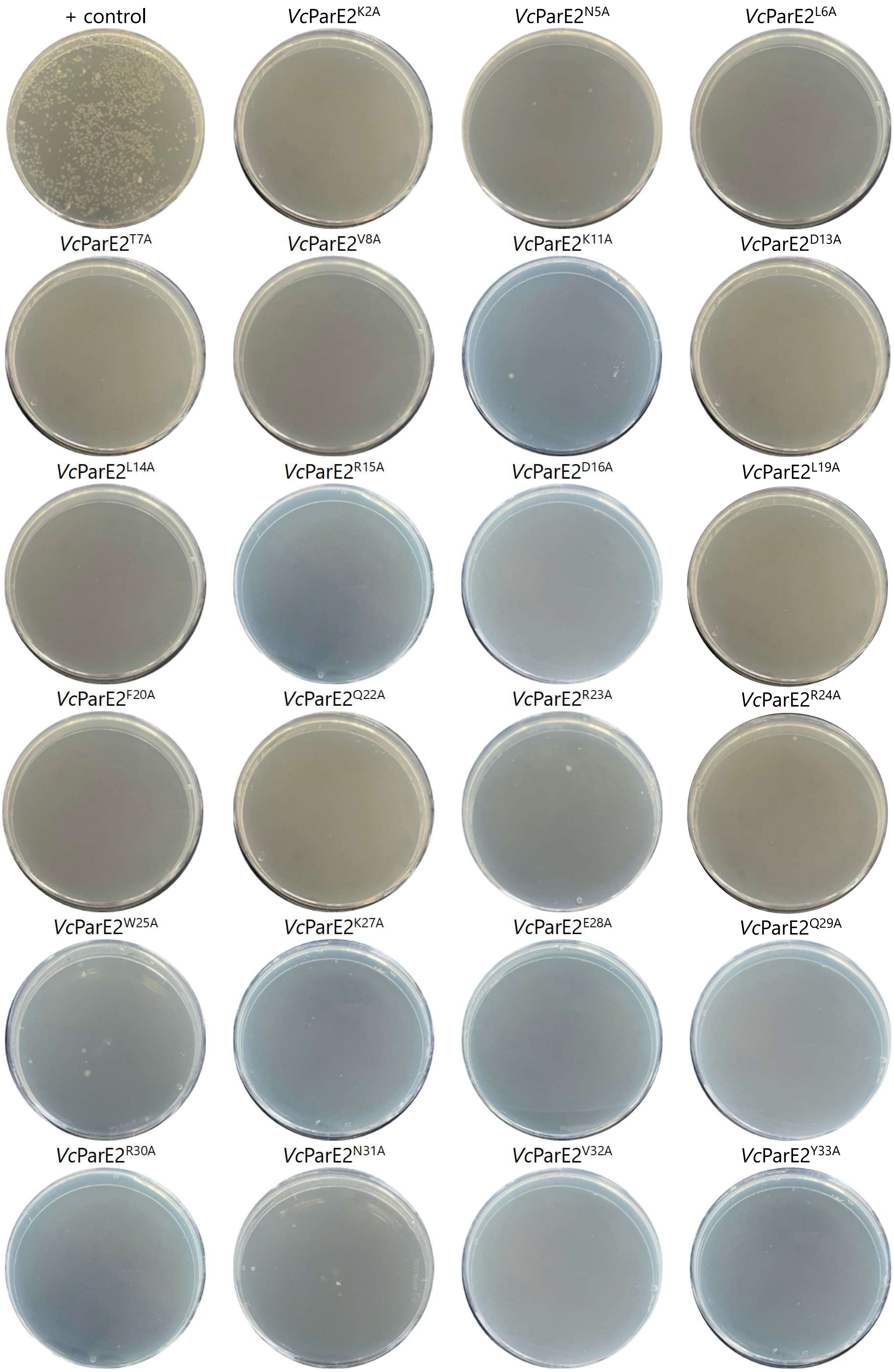
+ control
*Vc*ParE2$^{K2A}$
*Vc*ParE2$^{N5A}$
*Vc*ParE2$^{L6A}$
*Vc*ParE2$^{T7A}$
*Vc*ParE2$^{V8A}$
*Vc*ParE2$^{K11A}$
*Vc*ParE2$^{D13A}$
*Vc*ParE2$^{L14A}$
*Vc*ParE2$^{R15A}$
*Vc*ParE2$^{D16A}$
*Vc*ParE2$^{L19A}$
*Vc*ParE2$^{F20A}$
*Vc*ParE2$^{Q22A}$
*Vc*ParE2$^{R23A}$
*Vc*ParE2$^{R24A}$
*Vc*ParE2$^{W25A}$
*Vc*ParE2$^{K27A}$
*Vc*ParE2$^{E28A}$
*Vc*ParE2$^{Q29A}$
*Vc*ParE2$^{R30A}$
*Vc*ParE2$^{N31A}$
*Vc*ParE2$^{V32A}$
*Vc*ParE2$^{Y33A}$

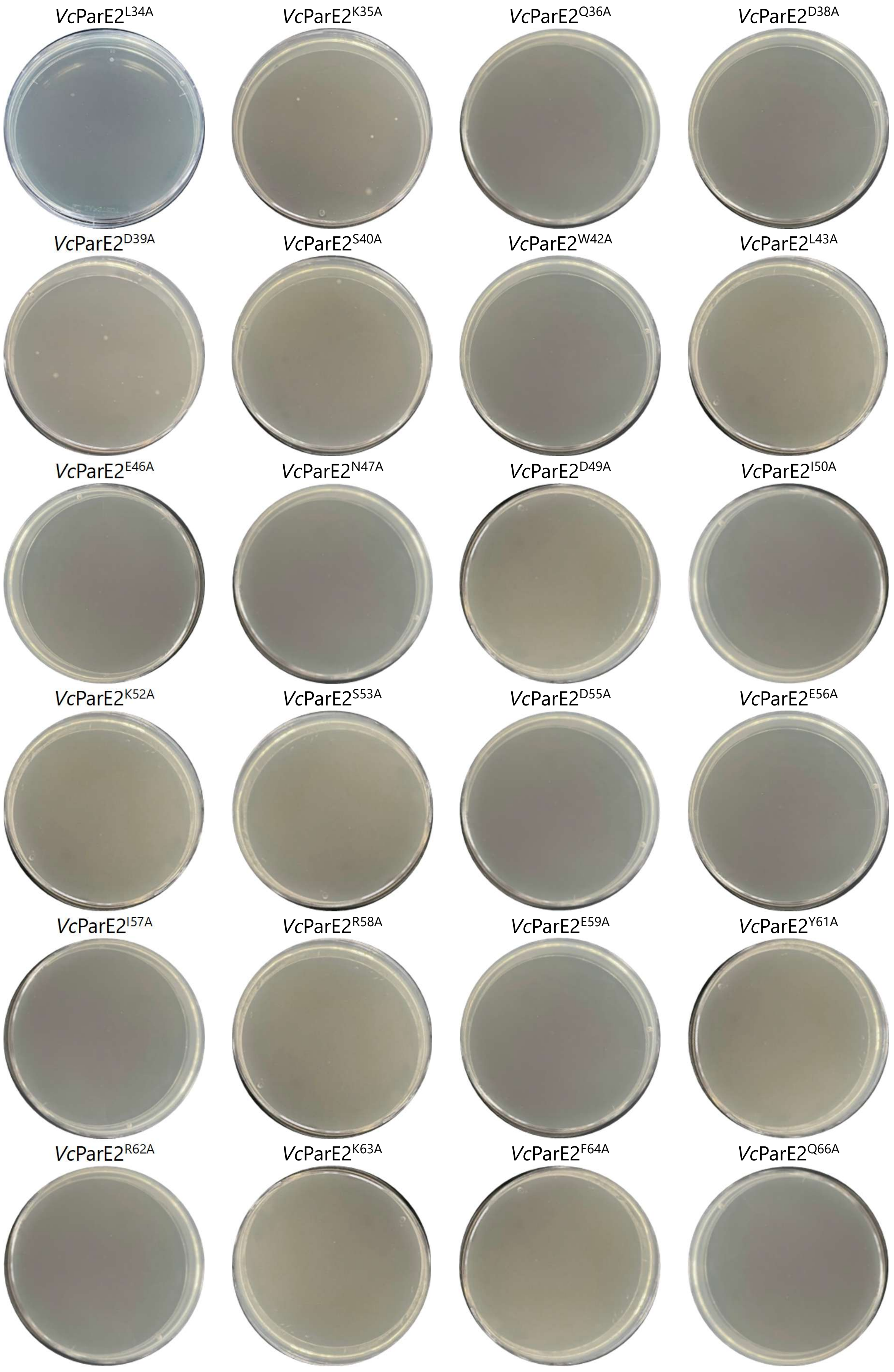
*Vc*ParE2$^{L34A}$
*Vc*ParE2$^{K35A}$
*Vc*ParE2$^{Q36A}$
*Vc*ParE2$^{D38A}$
*Vc*ParE2$^{D39A}$
*Vc*ParE2$^{S40A}$
*Vc*ParE2$^{W42A}$
*Vc*ParE2$^{L43A}$
*Vc*ParE2$^{E46A}$
*Vc*ParE2$^{N47A}$
*Vc*ParE2$^{D49A}$
*Vc*ParE2$^{I50A}$
*Vc*ParE2$^{K52A}$
*Vc*ParE2$^{S53A}$
*Vc*ParE2$^{D55A}$
*Vc*ParE2$^{E56A}$
*Vc*ParE2$^{I57A}$
*Vc*ParE2$^{R58A}$
*Vc*ParE2$^{E59A}$
*Vc*ParE2$^{Y61A}$
*Vc*ParE2$^{R62A}$
*Vc*ParE2$^{K63A}$
*Vc*ParE2$^{F64A}$
*Vc*ParE2$^{Q66A}$

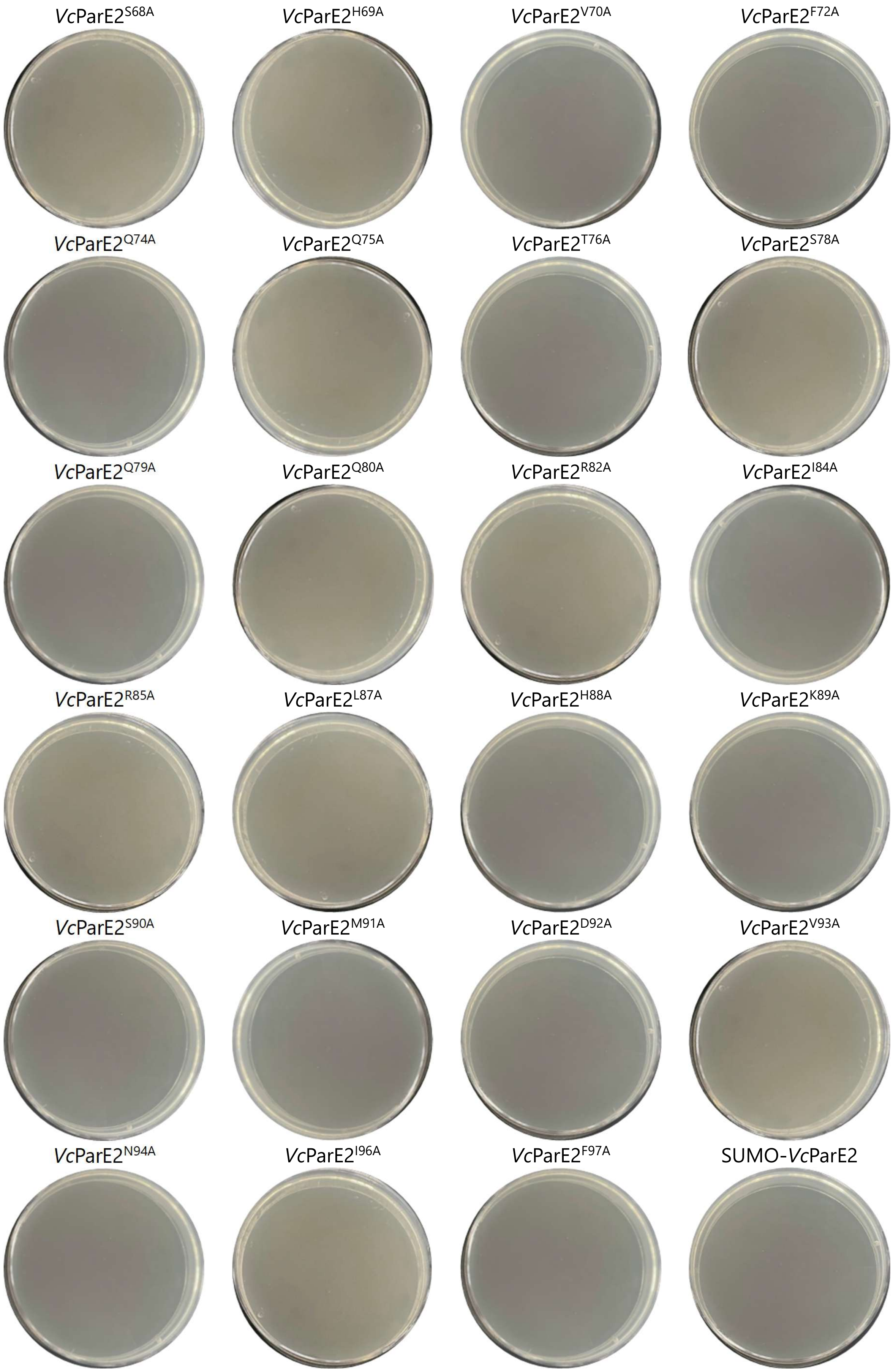
*Vc*ParE2$^{S68A}$
*Vc*ParE2$^{H69A}$
*Vc*ParE2$^{V70A}$
*Vc*ParE2$^{F72A}$
*Vc*ParE2$^{Q74A}$
*Vc*ParE2$^{Q75A}$
*Vc*ParE2$^{T76A}$
*Vc*ParE2$^{S78A}$
*Vc*ParE2$^{Q79A}$
*Vc*ParE2$^{Q80A}$
*Vc*ParE2$^{R82A}$
*Vc*ParE2$^{I84A}$
*Vc*ParE2$^{R85A}$
*Vc*ParE2$^{L87A}$
*Vc*ParE2$^{H88A}$
*Vc*ParE2$^{K89A}$
*Vc*ParE2$^{S90A}$
*Vc*ParE2$^{M91A}$
*Vc*ParE2$^{D92A}$
*Vc*ParE2$^{V93A}$
*Vc*ParE2$^{N94A}$
*Vc*ParE2$^{I96A}$
*Vc*ParE2$^{F97A}$
SUMO-*Vc*ParE2

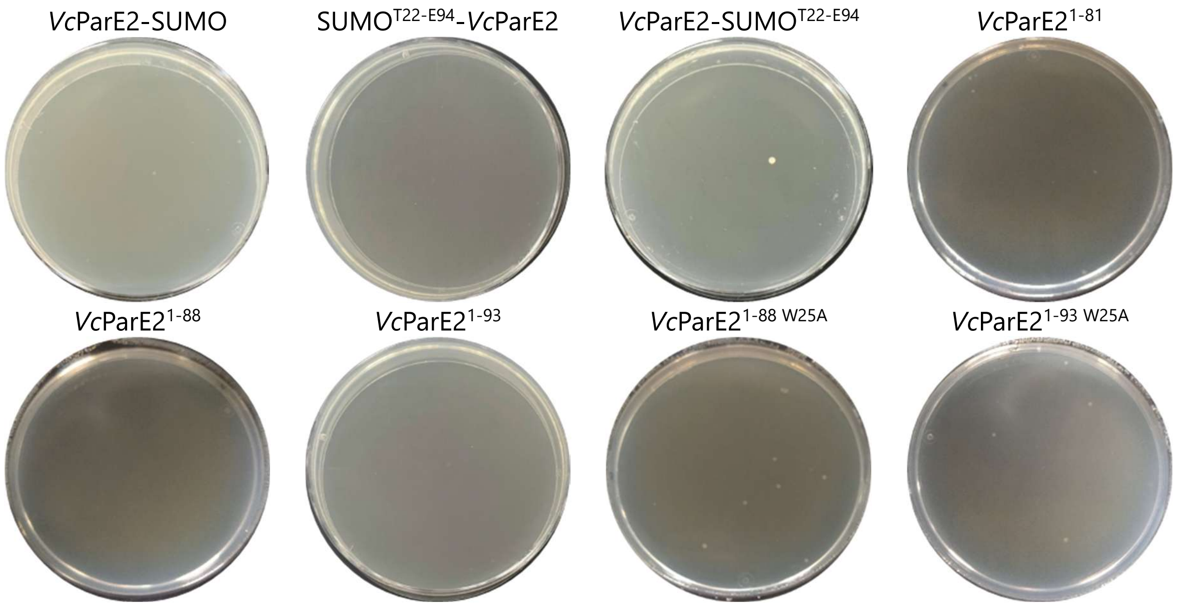


**Supplementary Figure S2:** Colony growth of *E. coli* NEB5α cells after transformation with pET28b-*Vc*ParE2mutant constructs. This figure presents an overview of all colonies that grew on LB plates supplemented with kanamycin following the transformation of *E. coli* NEB5α-cells with pET28b vectors encoding the indicated *Vc*ParE2 mutants. All colonies were screened using colony PCR and sequences were confirmed by Sanger sequencing. As a positive control, an empty pET28b vector was included. Fusions of *Vc*ParE2 and Small Ubiquitin-like Modifier (SUMO) protein have the full SUMO protein (PDB 1EUV, chain B) or only the folded segment of this protein (SUMO$^{T22-E94}$) attached at either the N-or C-terminus.

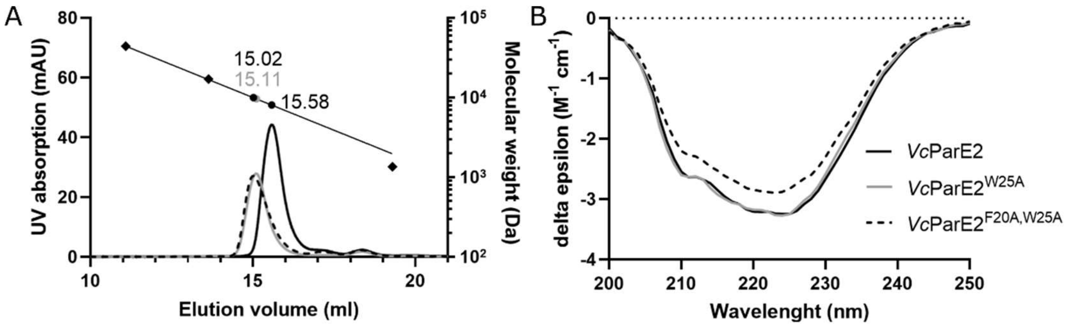


**Supplementary Figure S3:** *Vc*ParE2, *Vc*ParE2$^{W25A}$ and *Vc*ParE2$^{F20A,\ W25A}$ are well-folded monomers. (**A**) Analytical size exclusion chromatography profiles of wild type *Vc*ParE2 (solid black line), *Vc*ParE2 single mutant (solid grey line) and *Vc*ParE2 double mutant (dotted black line). Elution volumes of the BioRad molecular weight standards (chicken ovalbumin, 44 kDa; horse myoglobin, 17 kDa; vitamin B12, 1.35 kDa) are represented as diamonds and the regression curve is shown as a black line. (**B**) Circular dichroism profiles.

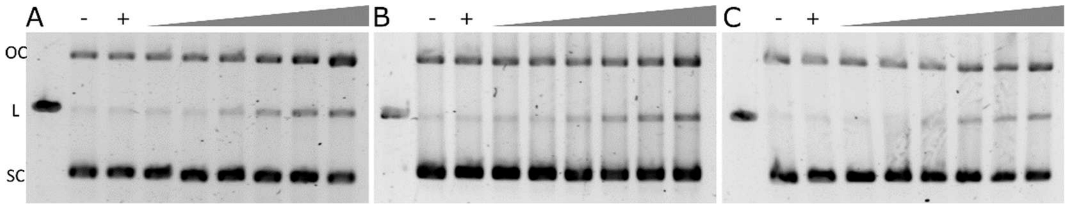


**Supplementary Figure S4:** Cleavage assays showing stabilisation of the cleavage complex by applying a gradient of 0.05, 0.1, 0.5, 1, 2.5 and 5 µM of (**A**) *Vc*ParE2$^{W25A}$, (**B**) *Vc*ParE2$^{F20A,\ W25A}$ and (**C**) moxifloxacin to a mixture of supercoiled pBR322 and Gyrase. The left lane in each gel corresponds with linearized pBR322 plasmid. DNA topoisomers are labelled as OC (open circular), L (linear) and SC (supercoiled). Negative control lanes (-) contain only negatively supercoiled pBR322 DNA, while positive control lanes (+) contain both DNA and Gyrase.

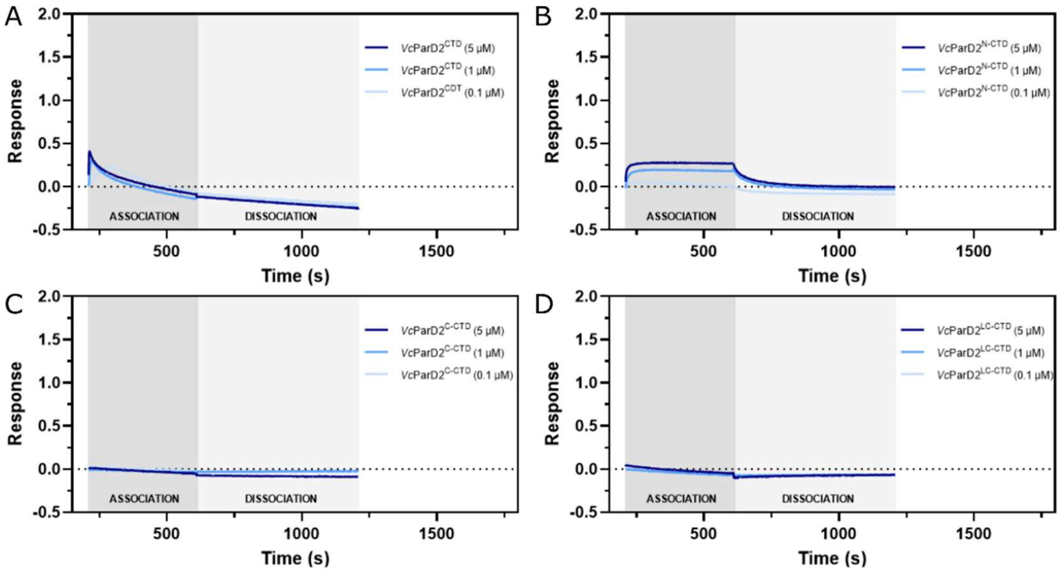


**Supplementary Figure S5:** BLI measurements of interactions between *Vc*ParE2 and different subfragments of the *Vc*ParD2 C-terminus. *Vc*ParE2 was coupled to the chip via its his-tag and submerged in a concentration series of the *Vc*ParD2 full C-terminus (residues E41-R80; **A**), the N-terminal helix of the *Vc*ParD2 C-terminus (residues E41-S60; **B**), the C-terminal helix of the *Vc*ParD2 C-terminus (residues D66-R80; **C**) or the loop and C-terminal helix of the *Vc*ParD2 C-terminus (residues G61-R80; **D**).
